# *NANOGP1*, a tandem duplicate of *NANOG*, exhibits partial functional conservation in human naïve pluripotent stem cells

**DOI:** 10.1101/2022.08.18.504441

**Authors:** Katsiaryna Maskalenka, Gökberk Alagöz, Felix Krueger, Joshua Wright, Maria Rostovskaya, Asif Nakhuda, Adam Bendall, Christel Krueger, Simon Walker, Aylwyn Scally, Peter J. Rugg-Gunn

**Author notes:** These authors contributed equally to the work (Katsiaryna Maskalenka, Gökberk Alagöz).

## Abstract

Gene duplication events are important drivers of evolution by providing genetic material for new gene functions. They also create opportunities for diverse developmental strategies to emerge between species. To study the contribution of duplicated genes to human early development, we examined the evolution and function of *NANOGP1*, a tandem duplicate of the key transcription factor *NANOG*. We found that *NANOGP1* and *NANOG* have overlapping but distinct expression profiles, with high *NANOGP1* expression restricted to early epiblast cells and naïve-state pluripotent stem cells. Sequence analysis and epitope-tagging of the endogenous locus revealed that *NANOGP1* is protein-coding with an intact homeobox domain. *NANOGP1* has been retained only in great apes, whereas Old World monkeys have disabled the gene in different ways including point mutations in the homeodomain. *NANOGP1* is a strong inducer of naïve pluripotency; however, unlike *NANOG*, it is not required to maintain the undifferentiated status of human naïve pluripotent cells. By retaining expression, sequence and partial functional conservation with its ancestral copy, *NANOGP1* exemplifies how gene duplication and subfunctionalisation can contribute to transcription factor activity in human pluripotency and development.

**Summary statement:** Establishing that *NANOGP1* has retained partial functional conservation with its ancestral copy *NANOG* sheds light on the role of gene duplication and subfunctionalisation in human pluripotency and development.

## INTRODUCTION

Gene duplication is an important driver of genome and species evolution. The majority of protein-coding genes and many non-coding regulatory sequences have arisen by duplication events (Magadum et al., 2013; Ohta, 2000). Most duplicated genes undergo functional decay due to silencing, loss-of-function mutations, or lack of required regulatory regions (Magadum et al., 2013). However, some duplicated genes are expressed, with the new copy either acquiring a novel function (neofunctionalisation) or sharing the ancestral function with the parental gene (subfunctionalisation). As a result, the emergence of a new copy of a gene or a regulatory sequence enables organisms to exploit new competitive advantages and to adapt to changing environments (Fares, 2014; Force et al., 1999; Kondrashov and Kondrashov, 2006).

Human evolution and development have been driven in many cases by the gain of low copy repeats called segmental duplications. Over 5% of the human genome consists of segmental duplications, typically with more than 90% identity shared between the ancestral and the duplicated copies (Bailey et al., 2002; Marques-Bonet et al., 2009a). This percentage of duplicated regions is remarkably high compared to Old World monkeys, such as macaques, where only 1.5% of the genome consists of such duplicates (Marques-Bonet et al., 2009a). A burst of duplication events followed the divergence of apes from Old World monkeys, and these copies account for ~80% of modern, human-specific duplications (Marques-Bonet et al., 2009b). For example, two gene duplicates –*SRGAP2C* and *ARHGAP11* – that are expressed in the developing human brain are proposed to have had a key role in the evolutionary expansion of the human neocortex (Charrier et al., 2012; Dennis and Eichler, 2016; Florio et al., 2015). However, the consequences of duplications underpinning such contributions remain largely undefined. Therefore, gene duplication events could be a major, unexplored driver of the divergence between mammalian developmental programmes yet, for most duplicated genes, their contribution to these early developmental programmes is poorly understood.

The core pluripotency transcription factor *NANOG* has a high number of duplicated copies in the human genome, and could therefore serve as a paradigm for studying the impact of gene duplication events on early development. High expression levels of *NANOG* are critical for maintaining the undifferentiated status of human naive and primed states of pluripotency (Guo et al., 2021; Hyslop et al., 2005; Lie et al., 2012; Vallier et al., 2009; Zaehres et al., 2005). If any of its duplicated copies are also highly expressed, that would raise the possibility that they might have an unanticipated role in human pluripotent cells. Ten of the eleven duplicates of *NANOG* are processed pseudogenes (copies of mRNAs that have been reverse transcribed and inserted into the genome), which lack regulatory sequences and possess various mutations that have led to their functional decay (Booth and Holland, 2004). Only one member of the *NANOG* pseudogene family – *NANOGP1* – is unprocessed (Booth and Holland, 2004). *NANOGP1* transcripts are detected in leukaemia cells, adult testes, and conventional or primed-state human pluripotent stem cells (hPSCs; naive-state hPSCs have not been examined) (Eberle et al., 2010; Hart et al., 2004). *NANOG* and *NANOGP1* share 97% coding region homology and have a similar exon-intron structure, suggesting that *NANOGP1* has probably undergone selection-driven conservation (Booth and Holland, 2004; Fairbanks and Maughan, 2006). Previous studies have reached contradictory conclusions about whether *NANOGP1* encodes a full-length protein (Booth and Holland, 2004; Eberle et al., 2010). If *NANOGP1* uses the equivalent translation initiation codon as *NANOG*,then, due to a base pair substitution, the resultant protein would contain only the first eight amino acid residues. However, *NANOGP1* could use an alternative, downstream initiation start codon that would encode a near full-length protein. This predicted NANOGP1 protein, if expressed, would have an intact homeodomain and transactivation domain, which are responsible for the protein dimerisation, DNA binding and pluripotency maintenance functions of *NANOG* and its orthologs (Chambers et al., 2003; Chang et al., 2009; Hart et al., 2004; Mullin et al., 2021; Oh et al., 2005; Theunissen et al., 2011). Whether endogenous *NANOGP1* can translate this protein has not been determined. This uncertainty about the predicted *NANOGP1* open reading frame led to the belief that *NANOGP1* does not encode a protein (Booth and Holland, 2004), and *NANOGP1* is currently classified as a non-protein-encoding pseudogene in the Ensembl repository.

Because NANOG has a central role in regulating human pluripotency, it is important to establish whether *NANOGP1* is a protein-coding gene that could also have functional capabilities. Here, we show that the NANOGP1 protein is expressed in naïve-state hPSCs. We determined that *NANOG* and *NANOGP1* have overlapping but not identical expression patterns in human embryos and stem cell lines. We found that, in contrast to *NANOG, NANOGP1* is not required to maintain undifferentiated naïve hPSCs, but *NANOGP1* can fulfil other functional roles of *NANOG* including reprogramming and autorepressive activities. By establishing that *NANOGP1* has retained partial functional conservation with its ancestral copy *NANOG*, our study sheds light on the role of gene duplication and subfunctionalisation on human pluripotency and development.

## RESULTS

### Identification of pseudogenes, including *NANOGP1*, that are highly expressed in human naïve pluripotent stem cells

To investigate pseudogene expression in human pluripotent cells, we first analysed transcript levels of pseudogenes in naïve-state hPSCs using RNA-sequencing. We selected 1,880 protein-coding genes in the human genome that have pseudogene copies (totalling 6,922 transcripts; Ensembl 104 annotation). Overall, 592 pseudogenes were detected with an expression value of log2RPM > 0 in naïve hPSCs (Fig. 1A). In particular, we found that several key pluripotency factors, including *NANOG, POU5F1* (also known as *OCT4*), and *DPPA3*, had highly expressed pseudogenes in naïve hPSCs (Fig. 1B, Fig. S1A-C). Four of these duplicated genes – *NANOGP1, POUF51P1, POU5F1P3* and *DPPA3P2* – were within the top 1% of all pseudogenes ranked by expression levels and their levels approached those of their ancestral copies (Fig. 1B). In addition to the duplicated pseudogene *NANOGP1* that was highly expressed, the processed and truncated genes *NANOGP4* and *NANOGP8* also had a substantial number of mapped reads (Fig. S1A). *POU5F1P1, POU5F1P3, DPPA3P2*,

**Figure S1.**
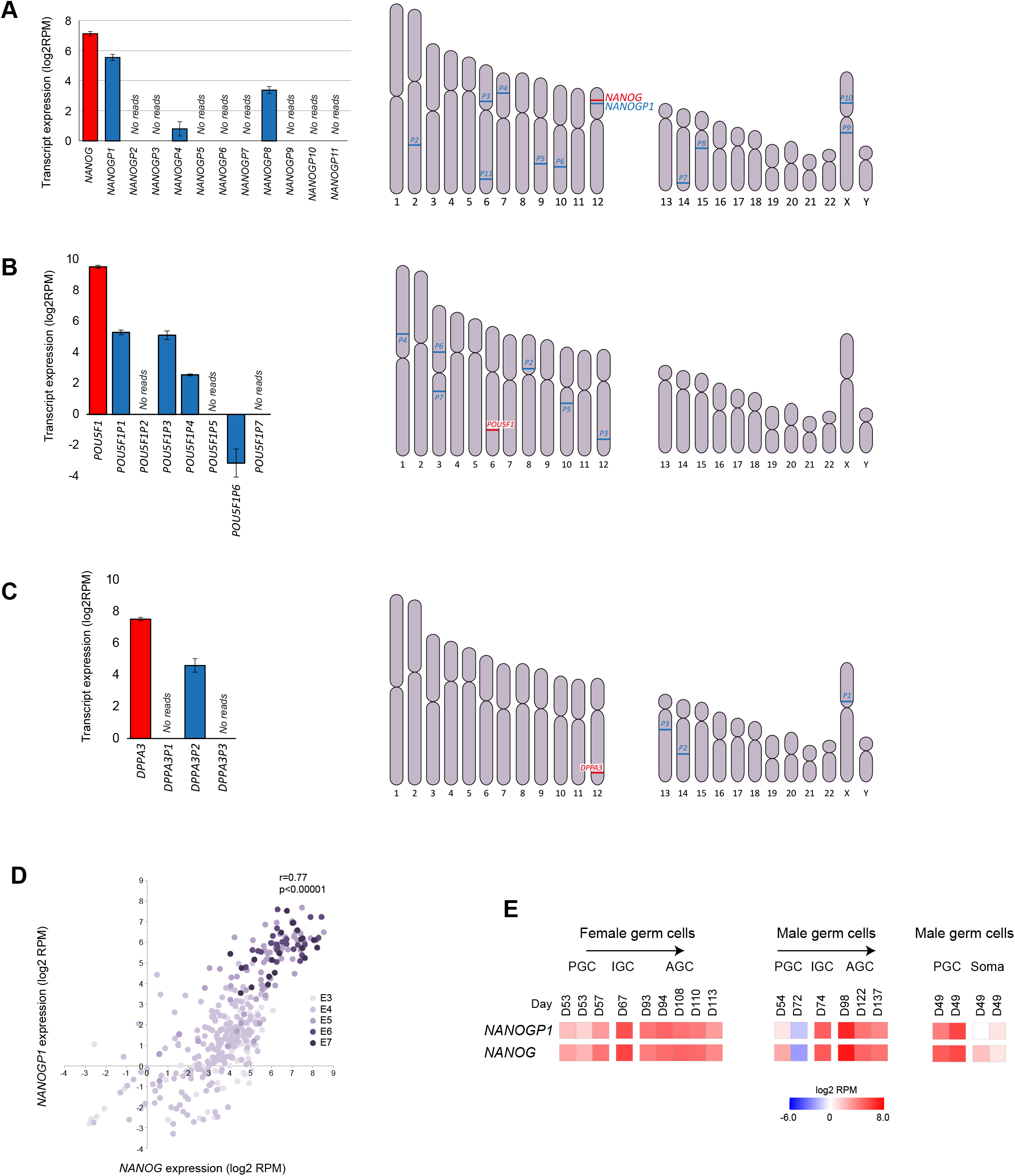
Overview of NANOG, POU5F1 and DPPA3 pseudogenes. **A–C)** *NANOG* (**A**), *POU5F1* (**B**) and *DPPA3* (**C**) transcript levels in naïve hPSCs (red) compared to the expression of their pseudogenes (yellow). Data show mean from three biologically independent samples ± SD. Idiograms show the chromosomal locations of *NANOG* (**A**), *POU5F1* (**B**) and *DPPA3* (**C**) and their pseudogenes. **D)** Scatter plot shows the expression of *NANOG* and *NANOGP1* expression in individual cells of the inner cell mass and epiblast lineages from embryonic day E3 to E7. Data were reanalysed from (Petropoulos et al., 2016). **E)** Heat maps show *NANOG* and *NANOGP1* expression in human male and female germ cells over the indicated days of foetal development. PGC, primordial germ cells; IGC, intermediate germ cells; AGC, advanced germ cells. Bulk RNA-seq data were re-analysed from (Gkountela et al., 2015).

*NANOGP4* and *NANOGP8* are processed copies, whereas *NANOGP1* was of specific interest because it has been formed by tandem duplication, is unprocessed, and is located in the same locus as its ancestral copy, *NANOG*. Together, these results uncover the large set of pseudogenes that are expressed in naïve hPSCs. In particular, the high expression of the duplicated pseudogene *NANOGP1* raises the possibility that this gene might have an unanticipated role in human pluripotent cells.

**Figure 1.**
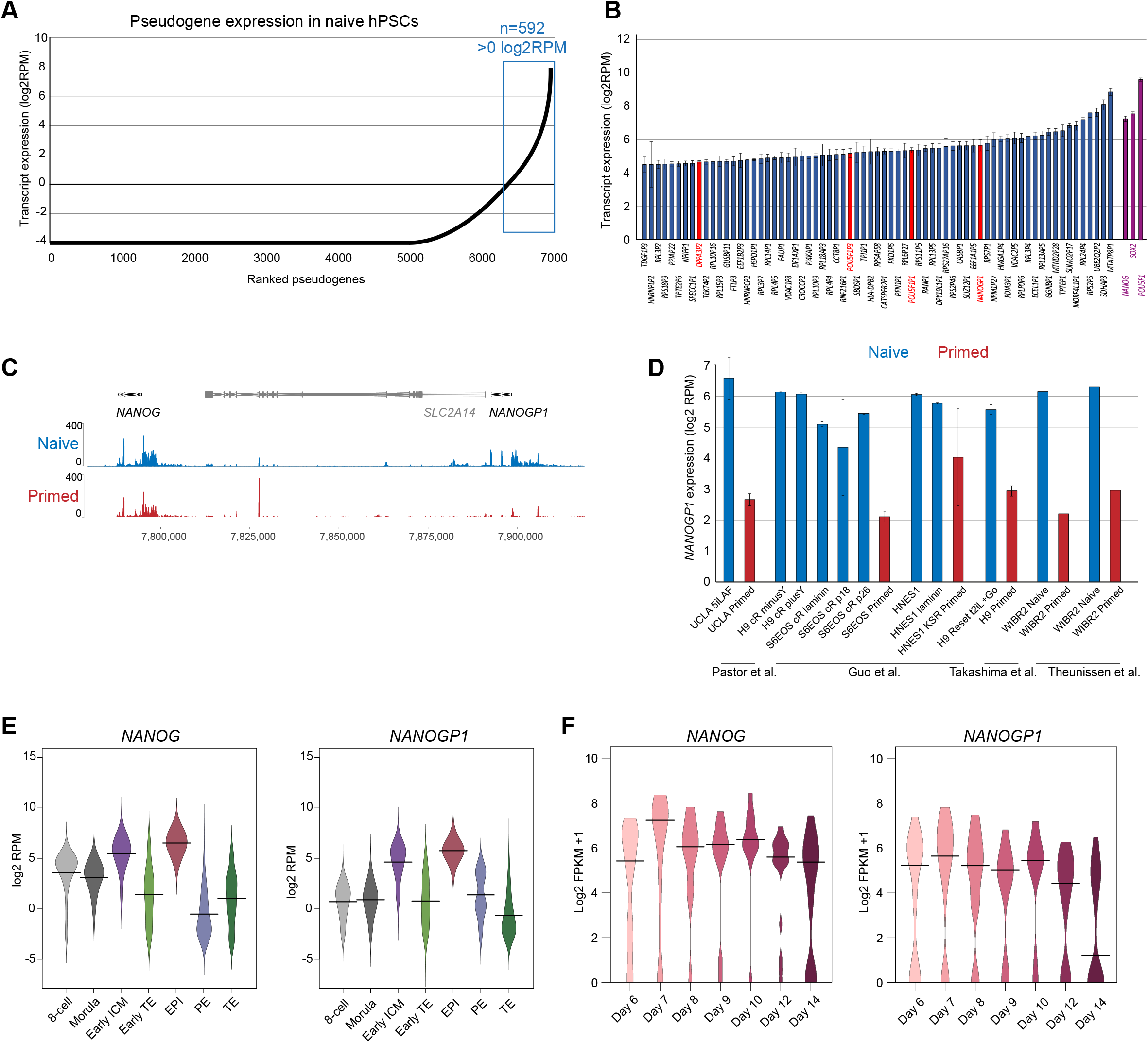
NANOGP1 is a highly expressed pseudogene in human naïve pluripotent stem cells and epiblast cells. **A)** Expression of 6,922 pseudogene transcripts in naïve hPSCs, ranked by expression level. **B)** Chart shows the top 1% (n=69) highest expressed pseudogenes in naïve hPSCs. Pseudogenes of pluripotency factors are highlighted in red. Three pluripotency factors – *NANOG, POU5F1* and *SOX2* – are shown for comparison. Data show mean from three biologically independent samples ± SD. **C)** Genome browser tracks of RNA-seq data for *NANOG, SLC2A14* and *NANOGP1* in naïve and primed hPSCs (H9 cell line). Data show merged tracks from three biologically independent samples (Collier et al., 2017). **D)** *NANOGP1* expression in multiple naïve (blue) and primed (red) hPSC lines. RNA-seq data was re-analysed from the indicated published studies (Guo et al., 2016; Pastor et al., 2016; Takashima et al., 2014; Theunissen et al., 2016), and includes naïve hPSCs generated by reprogramming and by direct derivation from blastocysts, and cultured in different conditions. For samples with error bars, the data show the mean from three biologically independent samples ± SD. **E)** *NANOG* and *NANOGP1* expression in human pre-implantation embryos in the indicated stages and lineages. 8 cell – 8-cell stage (n=78), Mor – morula (n=185), eICM – early inner cell mass (n=66), eTE – early trophectoderm (n=227), Epi - epiblast (n=45), PE – primitive endoderm (n=30), TE – trophectoderm (n=715). Horizontal line, median. Data were reanalysed from (Petropoulos et al., 2016). **F)** *NANOG* and *NANOGP1* expression in epiblast cells from human peri-implantation and early post-implantation cultured embryos across the indicated days. Day 6 (n=60); Day 7 (n=33); Day 8 (n=11); Day 9 (n=12); Day 10 (n=14); Day 12 (n=22); Day 14 (n=26). Horizontal line, median. Data were reanalysed from (Xiang et al., 2020).

### *NANOG* and *NANOGP1* have overlapping but distinct expression patterns

To study the expression pattern of *NANOGP1*, we next compared RNA-seq datasets of naïve and primed hPSCs, which are cell types that correspond to early and late epiblast cells of the human embryo, respectively. Although *NANOGP1* is a duplicated copy of *NANOG*, there were sufficient sequence differences between the transcripts of the two genes to uniquely assign RNA-seq reads to each gene (Sequence Divergence Rate of 0.013). We also confirmed that *NANOG* reads do not map to the *NANOGP1* locus and vice versa when using a high mapping quality value (MAPQ>20). The transcriptional analysis revealed notable differences in the expression patterns of *NANOG* and *NANOGP1*. Whereas *NANOG* is highly expressed in both naïve and primed hPSCs, *NANOGP1* is highly expressed only in naïve hPSCs and is substantially downregulated in primed hPSCs (Fig. 1C). Note that prior studies only examined primed hPSCs. This finding was confirmed and extended by analysing multiple RNA-seq data sets of different naïve and primed hPSC lines, including embryo-derived and reprogrammed cell lines, and cultured in different media conditions (Fig. 1D).

To test whether the distinct expression patterns are also observed *in vivo*, we reanalysed single-cell RNA-seq (scRNA-seq) data sets from human embryos (Petropoulos et al., 2016; Xiang et al., 2020). Like *NANOG, NANOGP1* was highly expressed in epiblast but not trophectoderm lineages (Fig. 1E). *NANOG* and *NANOGP1* expression was well-correlated in pre-implantation epiblast cells (Fig. S1E). Interestingly, we found that *NANOGP1* might be expressed in a subpopulation of primitive endoderm cells, although available cell numbers are low for this lineage (Fig. 1E). *NANOGP1* and *NANOG* transcripts were abundant throughout epiblast development, up until Day 14, at which point *NANOGP1* levels were abruptly reduced (Fig. 1F). In contrast, *NANOG* expression levels remained high including on Day 14 (Fig. 1F). This developmental expression pattern therefore mirrored the state-specific differences between naïve and primed hPSCs, further confirming the overlapping but distinct expression profiles of the two genes. Lastly, as *NANOG* is expressed in germ cells, we examined published RNA-seq data of *in vivo* germ cells (Gkountela et al., 2015) and found that *NANOGP1* transcripts are also detected at high levels that are comparable to *NANOG* (Fig. S1G). Overall, these results show that *NANOGP1* is dynamically expressed in hPSCs and developing human embryos, which is an expression pattern that is suggestive of a conserved potential role for *NANOGP1* in human early development.

### *NANOGP1* transcript and protein isoform sequences are highly similar to those of *NANOG*

The high expression and sequence read coverage of *NANOGP1* in naïve hPSCs enabled us to examine its mRNA structure, splicing patterns, and open reading frame sequences. This analysis identified three *NANOGP1* mRNA isoforms that differed due to alternative splicing between exons 3 and 4 (Fig. 2A). This pattern was consistent in additional naive hPSC lines from different studies (Fig. S2). No splicing to a putative upstream exon was detected, as had been previously considered (Booth and Holland, 2004). According to the splicing analysis in our study, the first *NANOGP1* exon was the same as that of *NANOG*. Due to a point mutation within exon 1, the most likely translation initiation codon for *NANOGP1* is 117 bp downstream of the equivalent initiation codon used by *NANOG* (Fig. 2B). This results in the open reading frame of NANOGP1 lacking the first 39 amino acids compared to NANOG (Fig. 2C), which is a finding that is consistent with earlier predictions (Booth and Holland, 2004; Hart et al., 2004). Outside of the first exon, the sequences encoding the main functional domains of NANOG, including the homeobox domain, tryptophan repeats and C-terminal transactivation domain, were all present and fully conserved in the predicted *NANOGP1* open reading frames (Fig. 2C). Several point mutations and two smaller deletions in isoforms 1 and 2 were detected outside of the main domains (Fig. 2C). Overall, these results show that the predicted sequences, exon structures and functional domains of *NANOGP1* are very similar to *NANOG*.

**Figure 2.**
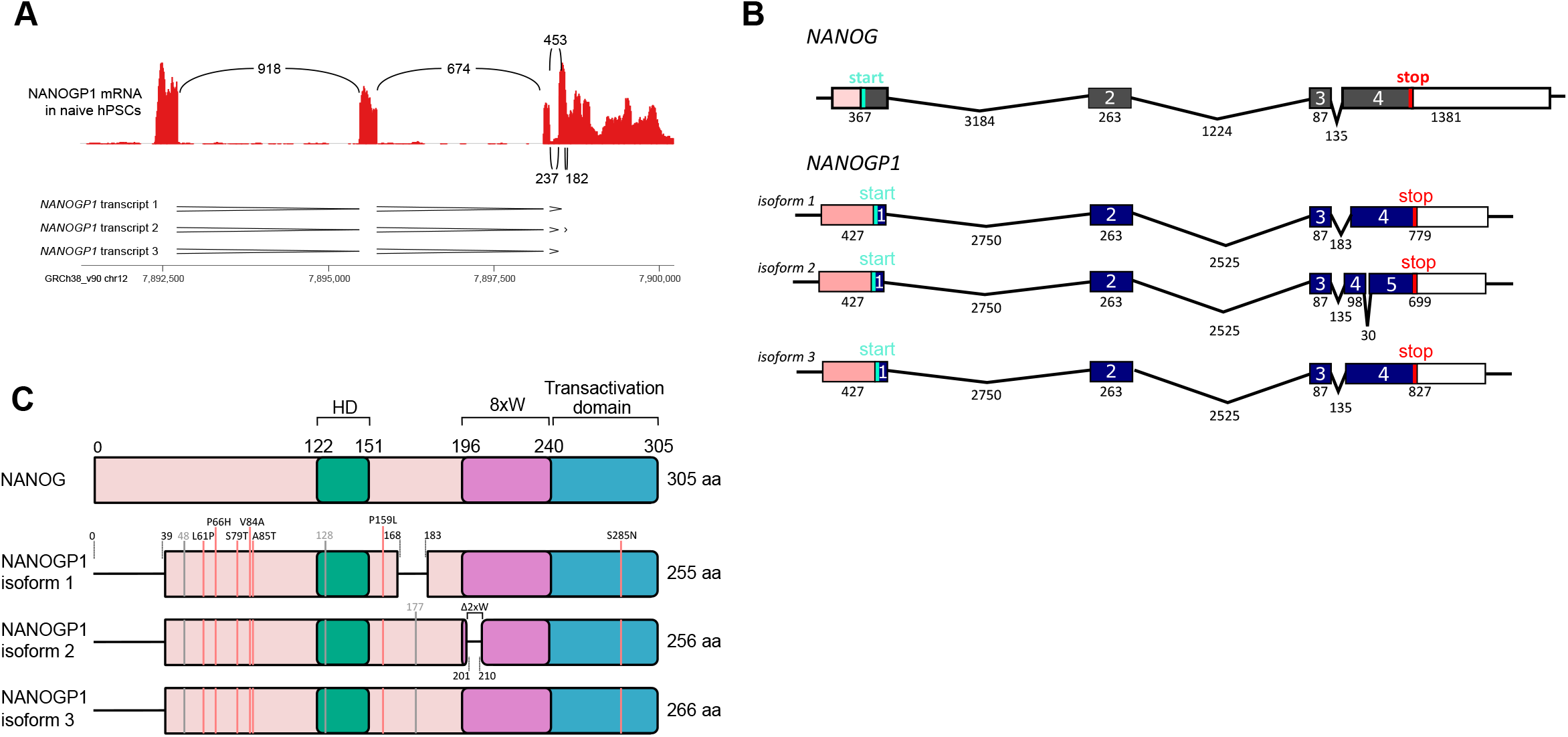
Splicing and sequence analyses reveal predicted open reading frame structure of NANOGP1. **A)** Sashimi plots show splicing analysis of *NANOGP1* transcripts in naïve hPSCs using RNA-seq data (Takashima et al., 2014). The numbers in between the RNA-seq peaks indicate the number of times a splicing event was measured. The three different predicted patterns of transcript splicing are indicated underneath. **B)** Schematic summarising the three predicted transcript isoforms of *NANOGP1*, including the size of each exon and intron (in bp) and translation start and start codons. *NANOG’s* transcript structure is shown for comparison. **C)** Diagram showing the three predicted NANOGP1 open reading frame (ORF) variants and domain structures based on the splicing and transcript analyses. The ORF of NANOG is shown for comparison. Differences in the NANOGP1 ORFs versus the NANOG ORF are indicated, including gaps. Amino acid substitutions caused by missense DNA changes are labelled by red vertical lines; silent changes are labelled by grey vertical lines. 8xW, tryptophan–rich subdomain/region containing 8 tryptophan (W) residues; Δ2xW, deletion of two tryptophan residues from the tryptophan-rich subdomain; HD, DNA-binding homeodomain.

### *NANOGP1* gene and protein sequences are highly conserved in Great Apes

We next examined the boundaries of the *NANOG/NANOGP1* duplication in the human genome. We self-aligned a 250 kb region containing *NANOG, NANOGP1, SLC2A14, SLC2A3*, and *NANOGNB*, plus their flanking regions on both sides (Fig. 3A). Three large domains of duplication were identified following this alignment: i) *NANOG* and *NANOGP1;* ii) *SLC2A14* and *SCL2A3;* and iii) an *SLC2A3* downstream region (Fig. 3A,B). These results are consistent with a duplication event that involved copying and inserting an ~80 kb region containing *NANOG* and *SLC2A14* into a new location immediately downstream of its original position, and which resulted in the formation of the *NANOG/NANOGP1* duplication.

**Figure 3.**
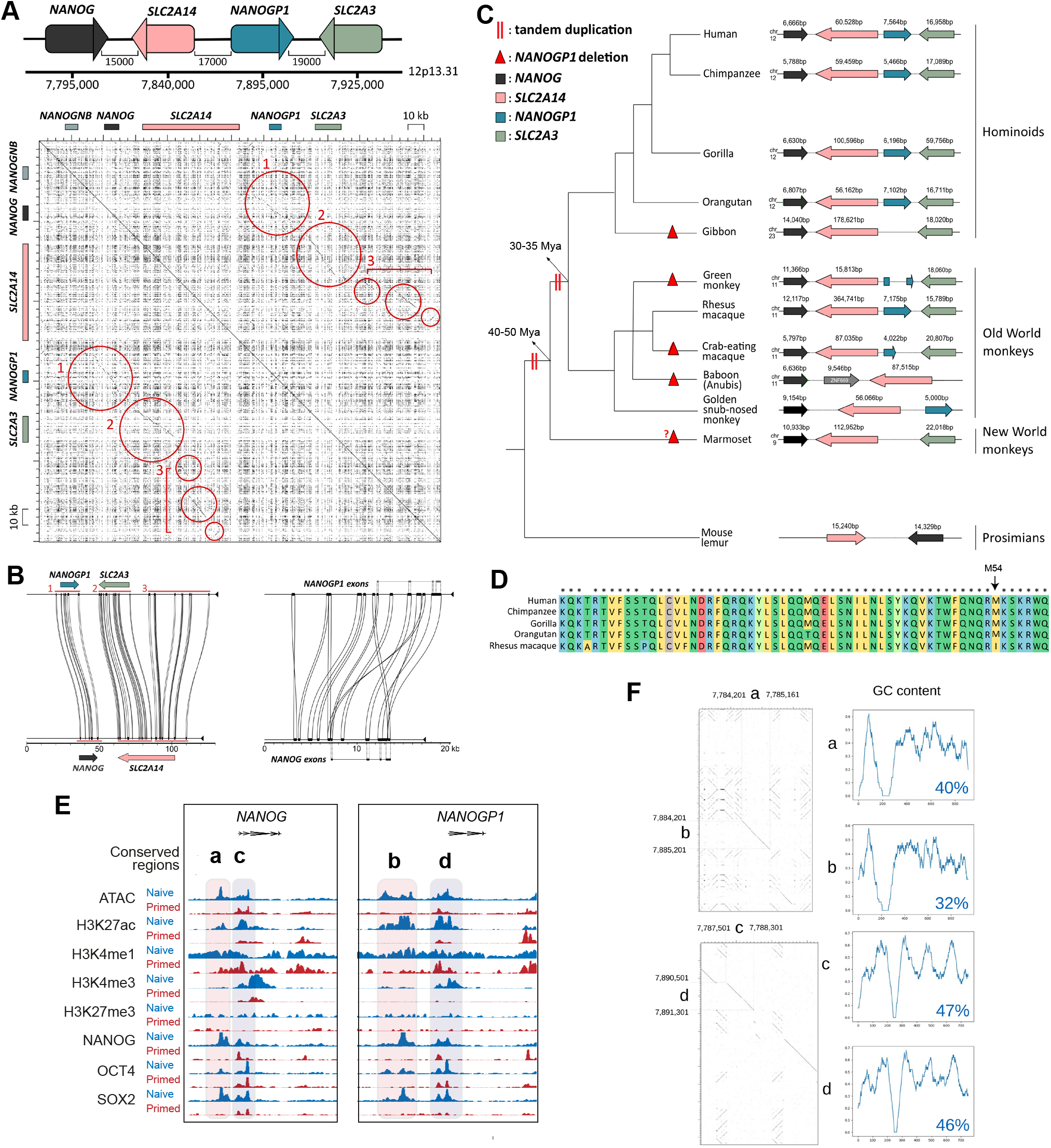
NANOGP1 duplication in human evolution. **A)** Top, diagram summarising the *NANOG/NANOGP1* tandem duplication locus (distance (bp) between the genes/pseudogene). Lower, dot plot shows self-alignment of a 250 kb region across the locus containing *NANOGNB, NANOG, NANOGP1* and another duplicated gene pair, *SLC2A14* and *SLC2A3* (genes indicated by boxes along x- and y-axes). Individual dots represent matching base pairs between the two aligned sequences. Circles indicate three areas of high sequence conservation between the ancestral and duplicated regions, which can be seen by the diagonal lines. **B)** Miropeats plots show sequence similarity and locations of the three regions identified in (**A**) (left) and between the exons and upstream regions of *NANOG* and *NANOGP1* (right). **C)** Conservation of the *NANOG/NANOGP1* tandem duplication locus across analysed species. Predicted duplication dates are indicated with two red vertical lines; predicted *NANOGP1* deletion events are indicated with red triangles. **D)** Amino acid alignment compares the homeodomain sequences of *NANOGP1* orthologs. Colour indicates different types of amino acids, according to their biochemical properties. *, amino acid is the same for all aligned sequences. **E)** Genome browser tracks of ATAC-seq (Pastor et al., 2016) and ChIP-seq (Chovanec et al., 2021) profiles across the *NANOG* and *NANOGP1* loci in naïve and primed hPSCs. The sequences labelled ‘a-d’ indicate two duplicated pairs of regulatory regions, with ‘a and b’ corresponding to putative enhancers, and ‘c and d’ representing promoters. **F)** Dot plots and GC content ratio line graphs showing comparison of the regulatory regions a-d. Individual dots represent matching base pairs between the two aligned sequences. In areas of sequence conservation individual dots form diagonal lines. GC content ratio graphs, where the x-axis represents the length of a putative regulatory region in bp, and the y-axis shows (G+C)/(G+C+A+T) values within sliding-windows of 30 bp. The average GC content ratios over the indicated regions are shown in the lower right corner of each graph.

To better understand the origins and conservation of the *NANOG/NANOGP1* duplication, we manually examined gene lengths, genomic positions and gene orientation data from genome assemblies of non-human apes, Old and New World monkeys and prosimians. We searched for unambiguous matches to *NANOGP1* in each assembly and annotated it where present, as this annotation was absent from most of the non-human genomes. We then aligned identified *NANOGP1* sequences to their corresponding *NANOG* counterparts (Fig. S3A,B). Our analysis revealed that the *NANOGP1* sequence is present in some ape and Old World monkey genomes, but not in New World monkey or prosimian genomes (Fig. 3C, Fig. S3A). This finding suggests that the duplication event occurred prior to the split between apes and Old World monkeys (30-35 million years ago, Mya) but more recently than the split between the Old World and New World monkeys (40-50 Mya) (Pozzi et al., 2014), and was followed by full or partial deletion on some lineages outside the

**Figure S2.**
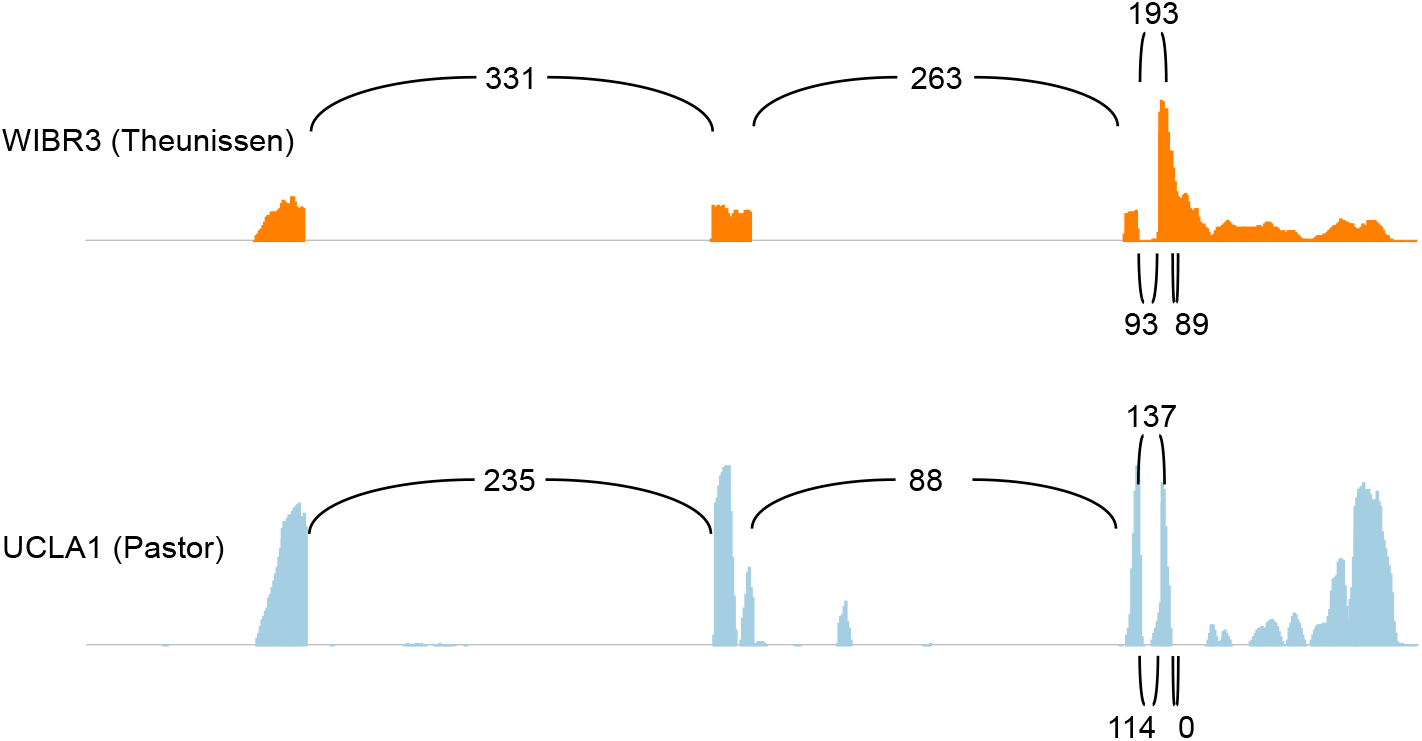
Examination of NANOGP1 in the genomes of non-human primates. Sashimi plots show splicing analysis of *NANOGP1* transcripts in naïve hPSCs using RNA-seq data from two additional studies using different cell lines (Pastor et al., 2016; Theunissen et al., 2016). The numbers in between the RNA-seq peaks indicate the number of times a splicing event was measured. All of the individual data sets examined revealed that there are three different predicted patterns of transcript splicing.

great apes (Fig. S3A-C). We note, however, that the marmoset genome (New World monkey) contains *SLC2A3*, which is a duplicated gene of *SLC2A14* (Fig. 3C). An alternative interpretation, therefore, is that the duplication event predated ~50 Mya and that *NANOGP1* was subsequently lost from the marmoset genome, or else that there were two separate duplication events: the first for *SLC2A14/SLC2A3* and the second for *NANOG/NANOGP1*.

*NANOGP1* sequences are present in most of the examined Old World monkey and ape species (Fig. 3C). Interestingly, however, an intact copy of *NANOGP1* is present only in great apes and, instead, the other species have inactivated *NANOGP1* in different ways. Some species, such as gibbon, have deleted the entire gene, whereas others, including the green monkey and crab-eating macaque, have partial deletions of *NANOGP1* (Fig. 3C, Fig. S3A-C). These species have retained *SLC2A3*. Other species appeared initially to have retained intact *NANOGP1*, but closer inspection uncovered small, critical mutations that are predicted to disable the protein. For example, *Rhesus macaque* contains a full-length *NANOGP1* sequence, but crucially has a non-synonymous amino acid change within the homeodomain (Fig. 3D). The affected amino acid, M54I, confers NANOG’s DNA binding specificity (Weiler et al., 1998). The likely consequence of this change is altered target sequence recognition because the homeobox protein PBX1, which also has an isoleucine at position 54, has a consensus motif of TGAT which differs from the canonical TAAT motif of NANOG (Chang et al., 1996; Piper et al., 1999). The function of *NANOGP1* in *Rhesus macaque* is therefore likely to be compromised. In contrast, the homeodomain sequences are intact for *NANOGP1* in human, chimpanzee and gorilla (Fig. 3D).

Taken together, these results show that a duplication event around 40 Mya created the *NANOG/NANOGP1* duplicated region that is present in the genomes of Old World monkeys and apes. *NANOGP1* has subsequently been disabled in most of the primate genomes via different alterations. Great apes, however, have retained intact gene and protein sequences, suggesting the potential presence of evolutionary pressure to maintain *NANOGP1* in those species.

### Putative regulatory regions upstream of *NANOGP1* were formed in the tandem duplication event

In addition to highly conserved exons, we also found distal regions that were conserved. Examining the sequence conservation and chromatin marks at the *NANOG/NANOGP1* locus revealed the location of several putative regulatory regions that overlapped with elements previously annotated as enhancers and super-enhancers (Fig. 3E and S4) (Chovanec et al., 2021). Six of these regions were identified near to *NANOGP1*, and four were positioned as two pairs directly upstream of *NANOG* (**a, c**) and *NANOGP1* (**b, d**) (Fig. 3E and S4). Pairwise alignments showed that the sequences within the two individual pairs, **a/b** and **c/d**, were very similar; additionally, each pair had matching GC content profiles, providing further evidence that they had formed from a duplication event (Fig. 3F). For the **c**/**d** pair, the GC content ratios were close to typical GC content ratio values that average ~50% in promoter regions (Villar et al., 2015), in contrast to the **a/b** pair that had lower GC content values (Fig. 3F). Together with the chromatin profiles, such as the promoter-

**Figure S3.**
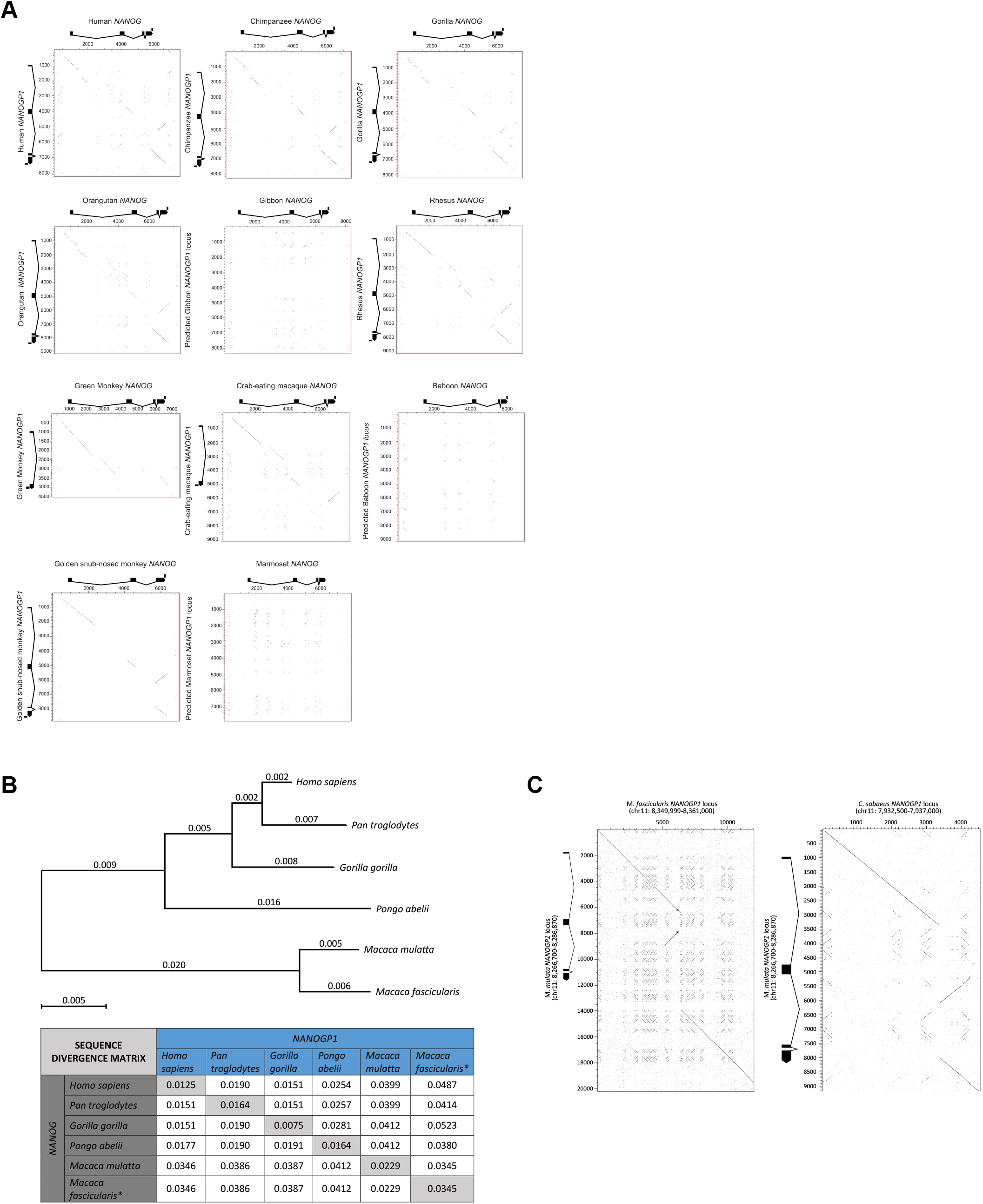
Examination of NANOGP1 in the genomes of non-human primates. **A)** Dot plots show the alignment of primate *NANOG* orthologs to their corresponding *NANOGP1* duplicates. Individual dots represent matching base pairs between the two aligned sequences. In areas of sequence conservation, individual dots form diagonal lines. Gene/pseudogene structure is shown as rectangles (exons) and lines (introns). Scale, bp. **B)** Upper, phylogenetic tree based on *NANOGP1* coding sequence. Neighbour-joining tree was based on the maximum likelihood model. Numbers on branches indicate evolutionary distance and correspond to substitutions/sequence length ratios. Substitutions are defined as nucleotides that are different from human *NANOGP1*. Lower, pairwise sequence divergence rates (# of substitutions/sequence length) of *NANOG* and *NANOGP1* coding sequences. Numbers correspond to substitutions per sequence length ratio. *In M. fascicularis genome, only 1st and 2nd exons are present. **C)** Dotter plots show partial *NANOGP1* deletions in green monkey and crab-eating macaque genomes.

**Figure S4.**
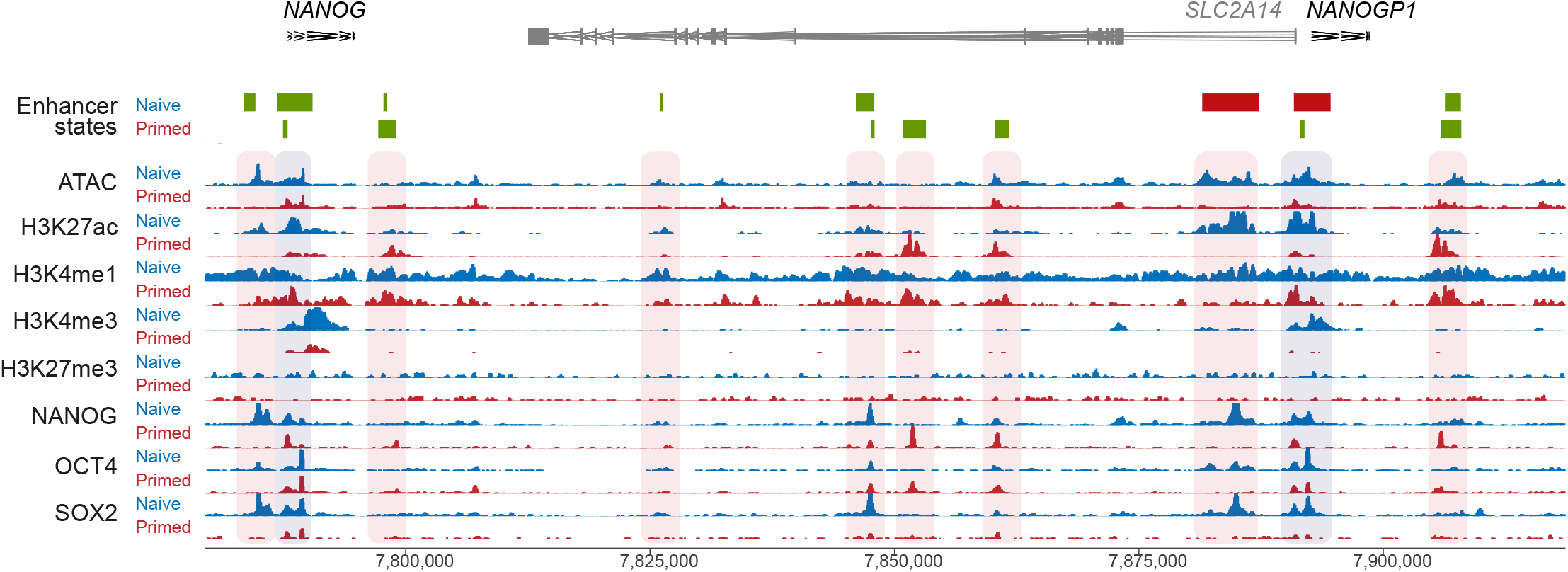
Characterisation of NANOGP1 putative regulatory sequences. Genome browser tracks of ATAC-seq (Pastor et al., 2016) and ChIP-seq (Chovanec et al., 2021) profiles across the *NANOG/NANOGP1* locus in naïve and primed hPSCs. The enhancer state tracks indicate the positions of previously defined enhancers (green boxes) and super-enhancers (red boxes) in each cell type; annotations from (Chovanec et al., 2021).

associated modification H3K4me3, this allowed us to conclude that **c/d** are likely to serve as promoters and **a/b** as enhancers.

According to ATAC-seq profiles (Pastor et al., 2018), sites **a, b, c** and **d** have highly accessible chromatin (Fig. 3E). Additionally, all four regions had high levels of active histone modifications – H3K27ac, H3K4me1 and H3K4me3 – and were bound by pluripotency factors in either one or both hPSC states (Fig. 3E) (Chovanec et al., 2021). The putative promoters **c** and **d** appeared active in both naïve and primed hPSC states and were hence referred to as ‘shared’, while the putative enhancers **a** and **b** were predominantly marked as active in the naïve hPSCs. The pattern of transcription factor occupancy and chromatin annotations were very similar for *NANOG* and *NANOGP1* at their putative promoter regions. The only prominent differences were for SOX2 and H3K4me3 levels within the shared putative promoters, where SOX2 and H3K4me3 peaks were detected near to *NANOG* in both primed and naïve hPSCs, but were present only in naïve hPSCs at the *NANOGP1* locus.

In summary, these results demonstrate that *NANOGP1* is integrated within the regulatory circuitry of pluripotent cells through OCT4, SOX2 and NANOG binding. The similarities in enhancer conservation and annotations could also help to explain the overlap of *NANOGP1* and *NANOG* expression patterns in human embryos and naïve hPSCs, and differences at the *NANOGP1* promoter in primed hPSCs correlate with reduced *NANOGP1* expression in those cells.

### *NANOGP1* encodes a protein that is expressed in naïve pluripotent stem cells

Although *NANOGP1* is currently annotated as a non-protein-encoding pseudogene, our revised sequence analysis suggested that the transcript should encode a protein of at least 255 amino acids. We therefore sought to establish whether NANOGP1 protein is detectable in naïve hPSCs. The close similarity in the predicted protein sequences of NANOGP1 and NANOG means there are no antibodies to detect NANOGP1 only. To overcome this, we used Cas12a ribonucleoprotein (RNP) and single stranded DNA (ssDNA) templates to insert V5 and 3xFLAG epitope tags into the endogenous *NANOGP1* coding sequence via homology directed repair (HDR) (Fig. 4A,B).

**Figure 4.**
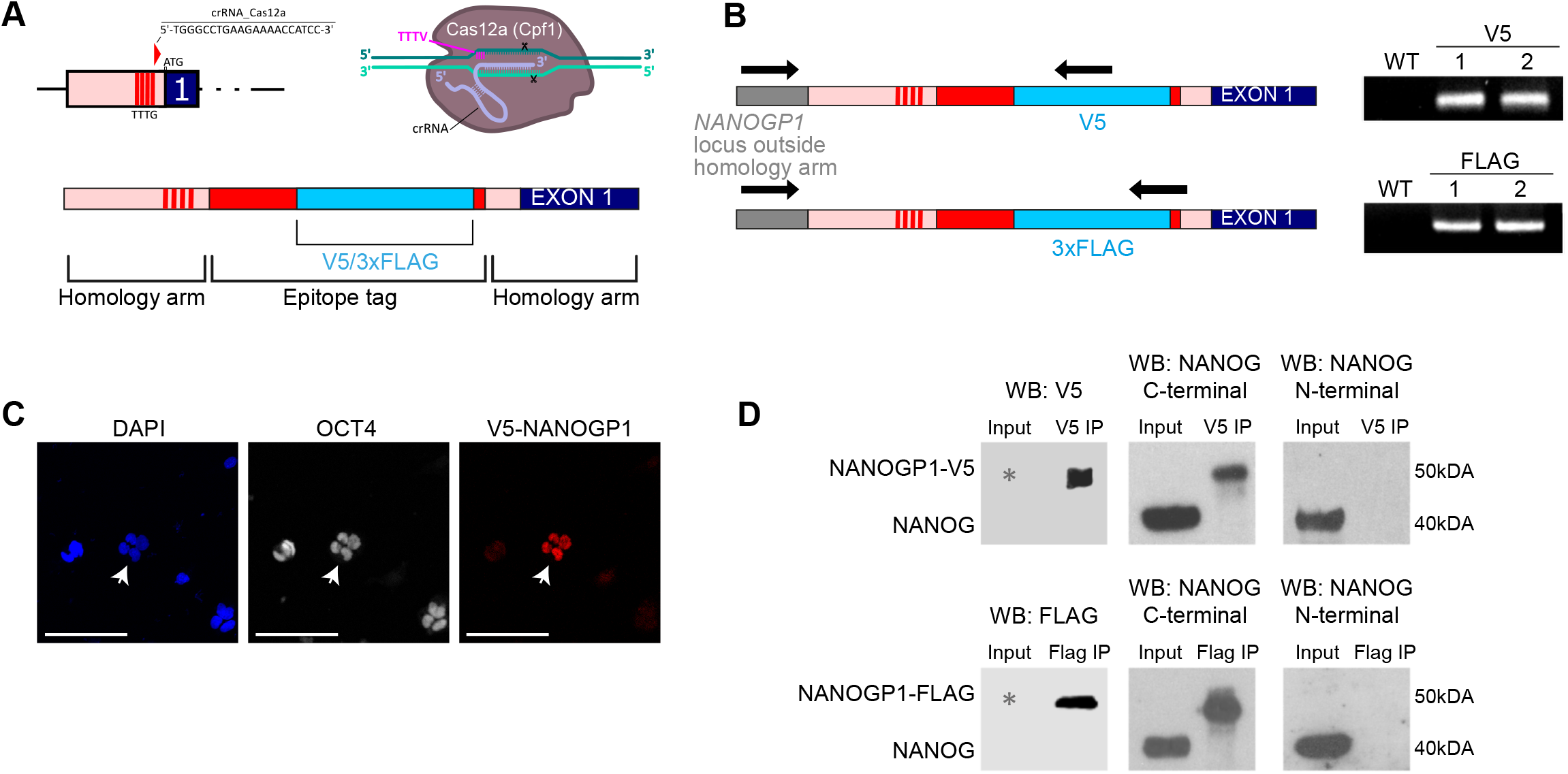
NANOGP1 encodes a protein that is expressed in human pluripotent cells. **A)** Schematic shows the CRISPR/Cas12a strategy to target the *NANOGP1* locus and insert an in-frame epitope tag. The crRNA recognises a sequence close to the *NANOGP1* translational start site. The single-stranded oligo DNA nucleotides used for homology-directed repair contains an in-frame sequence encoding either a V5 tag or a 3xFLAG tag, flanked by homology arms. **B)** Left, diagram shows the genotyping strategy where one primer (arrow) is at the *NANOGP1* locus outside of the homology arm, and the other primer (arrow) is within the epitope tag sequence. Right, PCR gel electrophoresis images confirm successful integration of the V5 and 3xFLAG tags into the *NANOGP1* locus in naïve hPSCs. WT, untransfected naïve hPSCs; V5-1 and V5-2, two independent naïve hPSC lines with V5 integrated at the *NANOGP1* locus; FLAG-1 and FLAG-2, two independent naïve hPSC lines with 3xFLAG integrated at the *NANOGP1* locus. **C)** Immunofluorescence microscopy images show nuclear localisation of V5-NANOGP1 in polyclonal transgenic naïve hPSCs, and overlap with OCT4 and DAPI signal. White arrows indicate the V5-positive colony. Scale bar, 100 μm. **D)** Western blot of co-immunoprecipitation experiments. Protein samples from transgenic polyclonal naïve hPSCs were immunoprecipitated with either V5 (upper) or FLAG (lower) antibodies. The immunoprecipitated material was examined by Western blot using antibodies against the epitope tag (left), the NANOG C-terminal that also detects NANOGP1 (centre), and the NANOG N-terminal that does not detect NANOGP1 due to an N-terminal deletion (right). The white asterisks indicate that due to the low number of NANOGP1-epitope tagged cells in the polyclonal population, the proteins were only detected in the immunoprecipitated samples and were not detected in the input samples.

We detected nuclear-localised expression of epitope-tagged NANOGP1 in polyclonal naïve hPSCs by immunostaining (Fig. 4C). Epitope-tagged NANOGP1 was also identified following immunoprecipitation and Western blotting (Fig. 4D). The specificity of the epitope-tagged protein was confirmed by using two different anti-NANOG antibodies for the Western blot: one that recognises the C-termini of NANOG and NANOGP1, and one that recognises the N-terminus of NANOG but not NANOGP1 (due to the N-terminal truncation of NANOGP1). These results establish that, in contrast to current annotations, *NANOGP1* is a protein-coding gene and its product is expressed in naïve hPSCs.

The discovery of NANOGP1 protein in naïve hPSCs prompted us to investigate whether this factor might have functional roles in naïve pluripotency. *NANOG* has several known functions in naïve pluripotent stem cells, including i) a gene autorepressive ability that was identified in mouse pluripotent stem cells (Navarro et al., 2012), ii) suppressing the transcription of the trophectoderm marker genes *GATA2, GATA3* and *TFAP2C* (Guo et al., 2021), and iii) reprogramming primed hPSCs towards the naïve state when overexpressed together with *KLF2* (Takashima et al., 2014; Theunissen et al., 2014). These three aspects of *NANOG* function were tested in relation to *NANOGP1* in the following sections.

### NANOGP1 has gene autorepressive activity

Ectopic *Nanog* overexpression in serum-free-cultured mouse pluripotent stem cells leads to the autorepression of endogenous *Nanog* expression by an unknown mechanism that likely involves NANOG binding upstream of its promoter (Navarro et al., 2012). To test whether *NANOG* and/or *NANOGP1* overexpression has a similar effect in human naïve pluripotency, we established hPSC lines containing doxycycline-inducible *NANOG* and *NANOGP1* transgenes (Fig. 5A,B). Transgenic naïve hPSCs were induced with doxycycline for 18 h and 72 h in t2iLGö media conditions (Fig. 5C,D). The induction of *NANOG* expression led to the downregulation of endogenous *NANOG* (Fig. 5C), thereby establishing that, as for mouse, human *NANOG* also has gene autorepressive activity. Interestingly, endogenous *NANOGP1* was also downregulated (Fig. 5C). Importantly, the overexpression of *NANOGP1* also suppressed the expression of *NANOG* and endogenous *NANOGP1* (Fig. 5D), thereby establishing that *NANOGP1* has a conserved autorepressive function.

**Figure 5.**
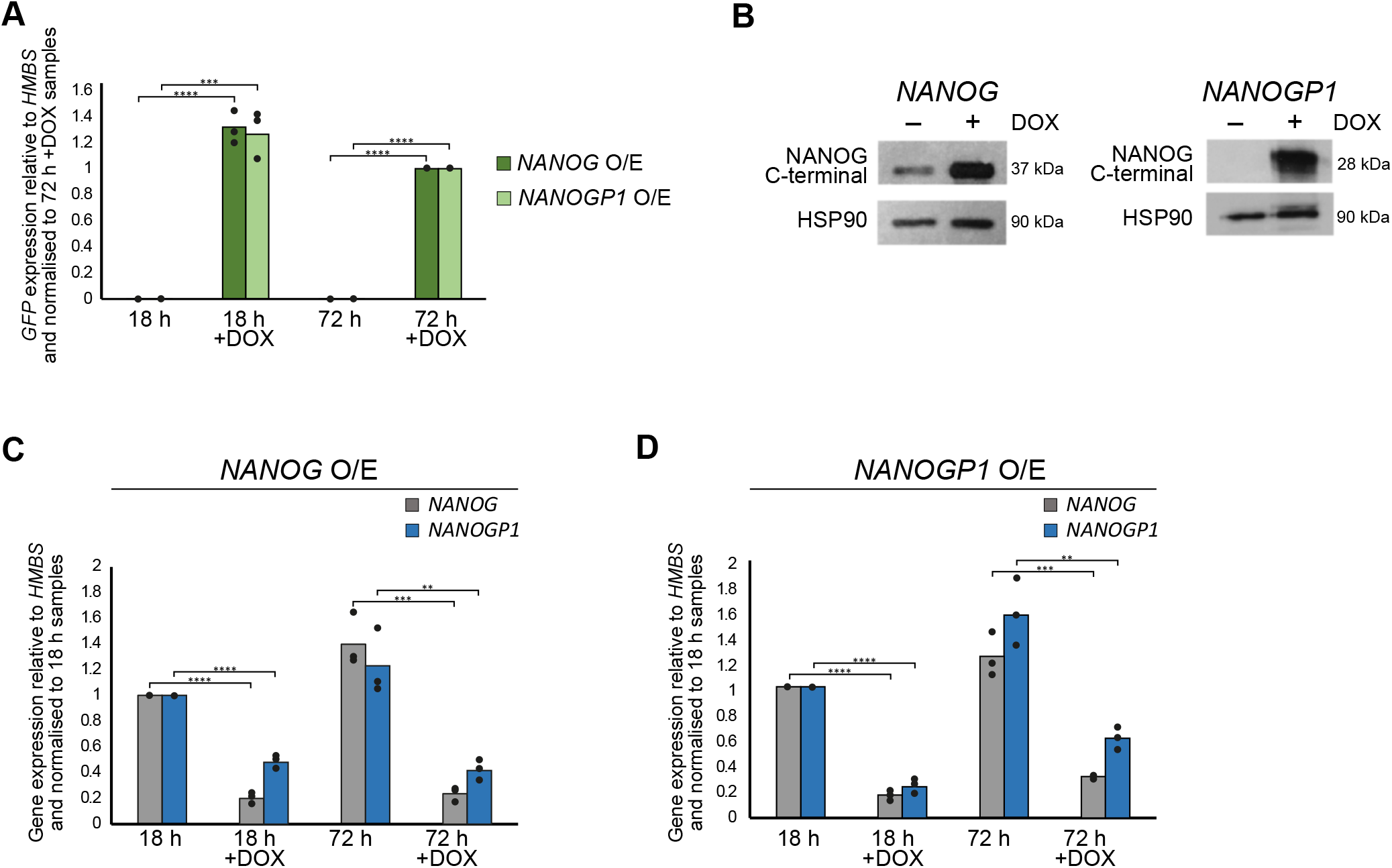
NANOGP1 has gene autorepressive activity. **A)** Induction of NANOG-GFP and NANOGP1-GFP transgenes in naïve hPSCs, as monitored by GFP expression. Naïve hPSCs were cultured in t2iLGo medium. RT-qPCR values are relative to HMBS expression and normalised to the 72 h + DOX samples. Mean and data points from three biologically independent samples are shown. Unpaired t-test (two-tailed) was performed (p = 0.0003 (***), p < 0.0001 (****)). **B)** Western blot showing DOX-induced overexpression of NANOG and NANOGP1 in naïve hPSCs. HSP90, loading control. **C** and **D)** Endogenous *NANOG* and *NANOGP1* expression levels in naïve hPSCs with DOX-inducible *NANOG* (C) and *NANOGP1* (D) transgenes. Primers target the 5’UTR of either *NANOG* or *NANOGP1*. RT-qPCR values are relative to *HMBS* expression and normalised to the 18 h samples. Mean and data points from three biologically independent samples are shown. Unpaired t-test (two-tailed) was performed (p < 0.01 (**), p < 0.001 (***), p < 0.0001 (****)).

### *NANOGP1* can reprogramme human primed pluripotent stem cells into a naïve state

The short-term, enforced expression of *NANOG* and *KLF2* facilitates the reprogramming of primed hPSCs into the naïve state (Takashima et al., 2014; Theunissen et al., 2014). We therefore investigated whether *NANOGP1* is also capable of promoting primed to naïve reprogramming, to ascertain whether *NANOGP1* can fulfil the role of *NANOG* in a direct functional test. *NANOGP1* was overexpressed together with *KLF2* in primed hPSCs using a doxycycline-inducible system in minimal 2i+LIF medium (Fig. 6A). We tested all three *NANOGP1* isoforms separately. To monitor and select for transgene expression, *NANOGP1* was co-translated with *GFP* via an internal ribosome entry site, and *KLF2* with *RFP*. Prior to reprogramming, we ensured comparable overexpression levels in all lines by inducing the cells with doxycycline for 24 h and flow-sorted the appropriate GFP+RFP+ or RFP+ only cell populations (Fig. S5A). The following day, the cells were switched to 2i+LIF medium with doxycycline to initiate reprogramming.

**Figure 6.**
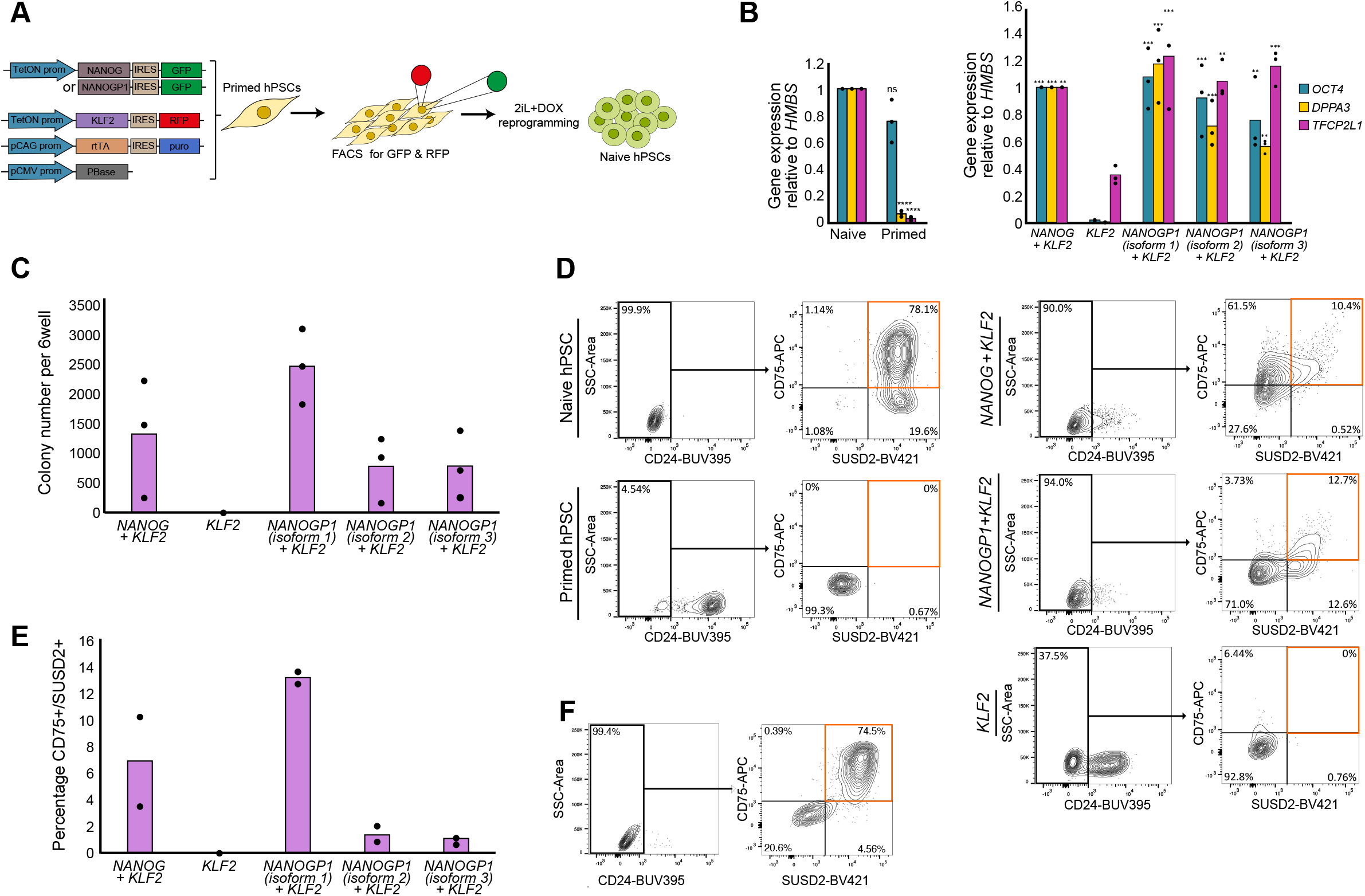
NANOGP1 is a strong inducer of naïve pluripotency. **A)** Schematic of experimental design for transgene-induced primed to naïve hPSC reprogramming. Plasmids encoding DOX-inducible *NANOGP1-ires-GFP* or *NANOG-ires-GFP, KLF2-ires-RFP*, and *pCAG-rtTA* and *pCMV-PBase*, were co-transfected into primed hPSCs. After a short pulse of DOX, GFP and RFP double positive cells were isolated by flow sorting, and transferred into 2iLIF medium supplemented with DOX. **B)** Expression of pluripotency markers in established naïve and primed hPSCs (left) and in cultures after 12 days of DOX-induced reprogramming (right). RT-qPCR values are relative to *HMBS* expression and normalised to naïve hPSCs (left) and to the *NANOG+KLF2* sample (right). All three *NANOGP1* isoforms were tested. Mean and data points from three biologically independent experiments are shown. One-way ANOVA with Dunnett’s multiple comparisons test compared all samples to the *KLF2*-only sample (p < 0.05 (*), p < 0.005 (**), 0.0005 (***), p < 0.00005 (****)); right) and t-test compared the primed sample to the naive samples (ns – not significant, p < 0.00005 (****); left). **C)** Chart showing the number of alkaline phosphatase-positive colonies after 12 days of DOX-induced reprogramming. Mean and data points from three independent reprogramming experiments are shown. **D)** Flow cytometry contour plots of cell-surface marker expression in established naïve and primed hPSCs (blue shading) and in cultures after 12 days of DOX-induced reprogramming (red shading). Naïve hPSCs (CD24 negative; CD75 positive; SUSD2 positive) are shown in the upper right quadrant of the final gate. **E)** Summary of the flow cytometry data from (**D**) for two independent reprogramming experiments. **F)** Flow cytometry contour plots confirming stable cell-surface marker expression in established *NANOGP1+KLF2* (isoform 1) cell lines propagated in the absence of DOX in naïve hPSC medium for 7 passages.

By Day 12 of reprogramming in these conditions, we observed numerous domed colonies with naïve hPSC morphology in the *NANOGP1+KLF2* cultures. The cells had upregulated naïve pluripotency markers, including *DPPA3* and *TFCP2L1*, and maintained high *POU5F1* expression (Fig. 6B). All three *NANOGP1* isoforms showed similar effects. These changes were comparable to the positive control cells expressing *NANOG* and *KLF2*. The reprogrammed colonies were positive for alkaline phosphatase activity, and the number of positive

**Figure S5.**
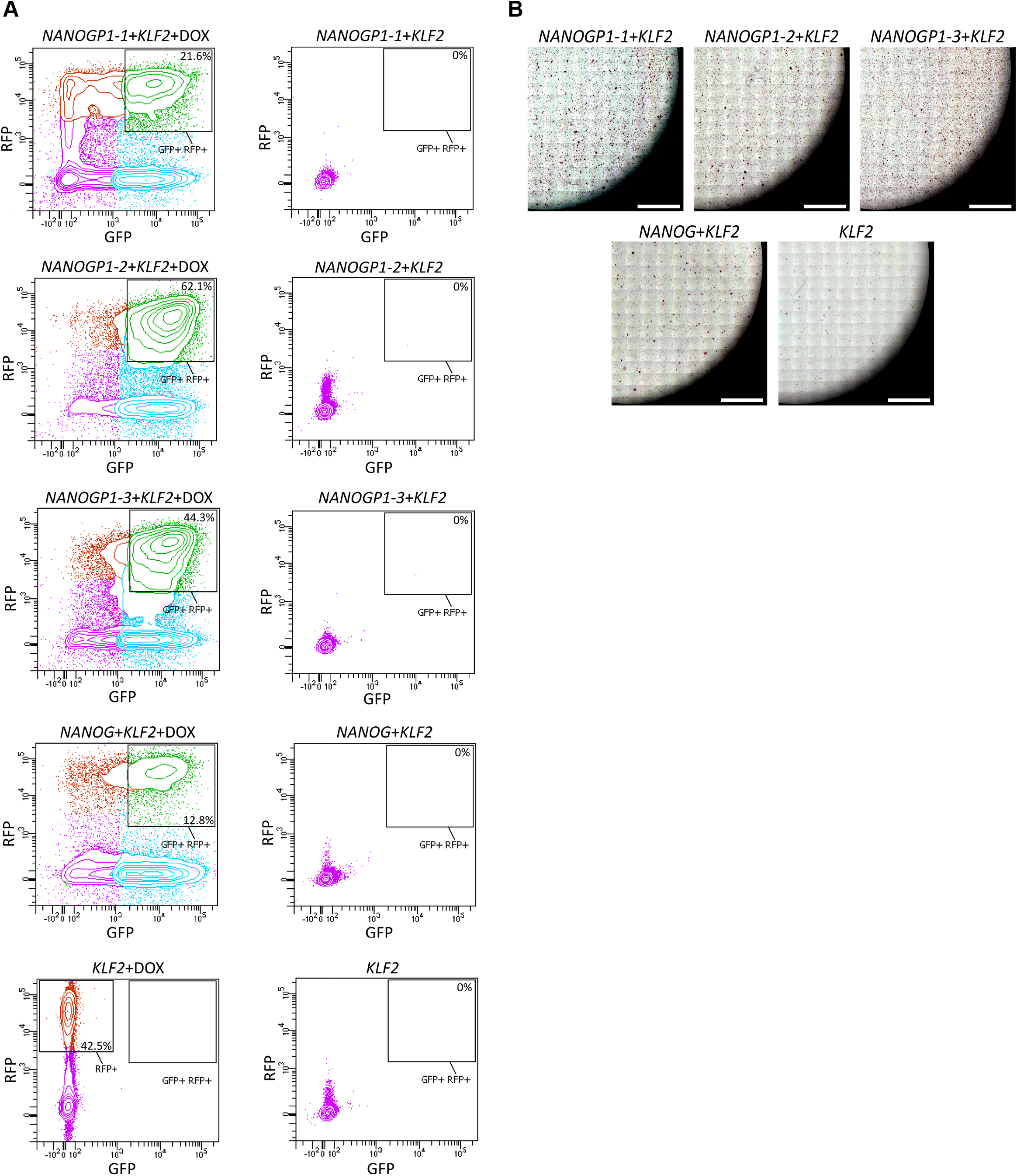
Characterisation of transgene-induced primed to naïve hPSC reprogramming. **A)** Flow cytometry contour plots show RFP and GFP expression in transgenic primed hPSCs. Samples treated with DOX for 48 h are shown on the left; non-treated samples on the right. Percentages of GFP+RFP+ and RFP+ populations are indicated. Data are representative of three biologically independent experiments. **B)** Brightfield microscopy images of the alkaline phosphatase assay. Reprogrammed naïve hPSC colonies are stained in purple. Scale, 5 mm.

colonies was similar when comparing cultures overexpressing either *NANOGP1* or *NANOG* (Fig. 6C, S5B). Flow cytometry analysis using stringent cell-surface markers of naïve pluripotency (CD24 negative; CD75 positive; SUSD2 positive) (Bredenkamp et al., 2019a; Collier et al., 2017; Shakiba et al., 2015; Wojdyla et al., 2020) validated successful pluripotent state conversion in the *NANOGP1*-overexpressing cells (Fig. 6D,E). Importantly, in all of the assays, the overexpression of *KLF2* alone did not induce reprogramming, confirming the critical contribution of *NANOGP1* in establishing naïve pluripotency. The change in pluripotent state was stable because the *NANOGP1*-induced reprogrammed cells retained their cell-surface marker phenotype when cultured for seven passages without doxycycline (Fig. 6F). Overall, these results lead us to conclude that, like *NANOG, NANOGP1* is capable of reprogramming hPSCs into the naïve state, thereby demonstrating functional conservation in igniting the naïve pluripotency network.

### *NANOGP1* is not required to maintain naïve pluripotency, unlike *NANOG*

We next set out to investigate whether *NANOGP1* supports the maintenance of human naïve pluripotency. A recent study showed that polyclonal cultures of *NANOG*-deficient naïve hPSCs upregulate several trophectoderm lineage marker genes, thereby uncovering a potentially crucial role for *NANOG* in maintaining naïve pluripotency (Guo et al., 2021). However, the dynamics of the transcriptional response following *NANOG* perturbation, and the effect on gene expression programmes, has not been examined. We first aimed at better defining this important phenotype, which would also provide a suitable comparison for studying whether the loss of *NANOGP1* might show similar effects.

We established naïve hPSC lines expressing doxycycline-inducible CRISPRi (dCas9-KRAB) (Mandegar et al., 2016) that targeted the promoters of either *NANOG* or *NANOGP1* by gene-specific gRNAs (Fig. 7A). Treating the transgenic naïve hPSC lines with doxycycline in t2iLGö medium caused the efficient and gene-specific knockdown of *NANOG* transcripts by 80%, and *NANOGP1* levels by 90% (Fig. 7B). NANOG protein was also strongly reduced after doxycycline treatment (Fig. 7C).

**Figure 7.**
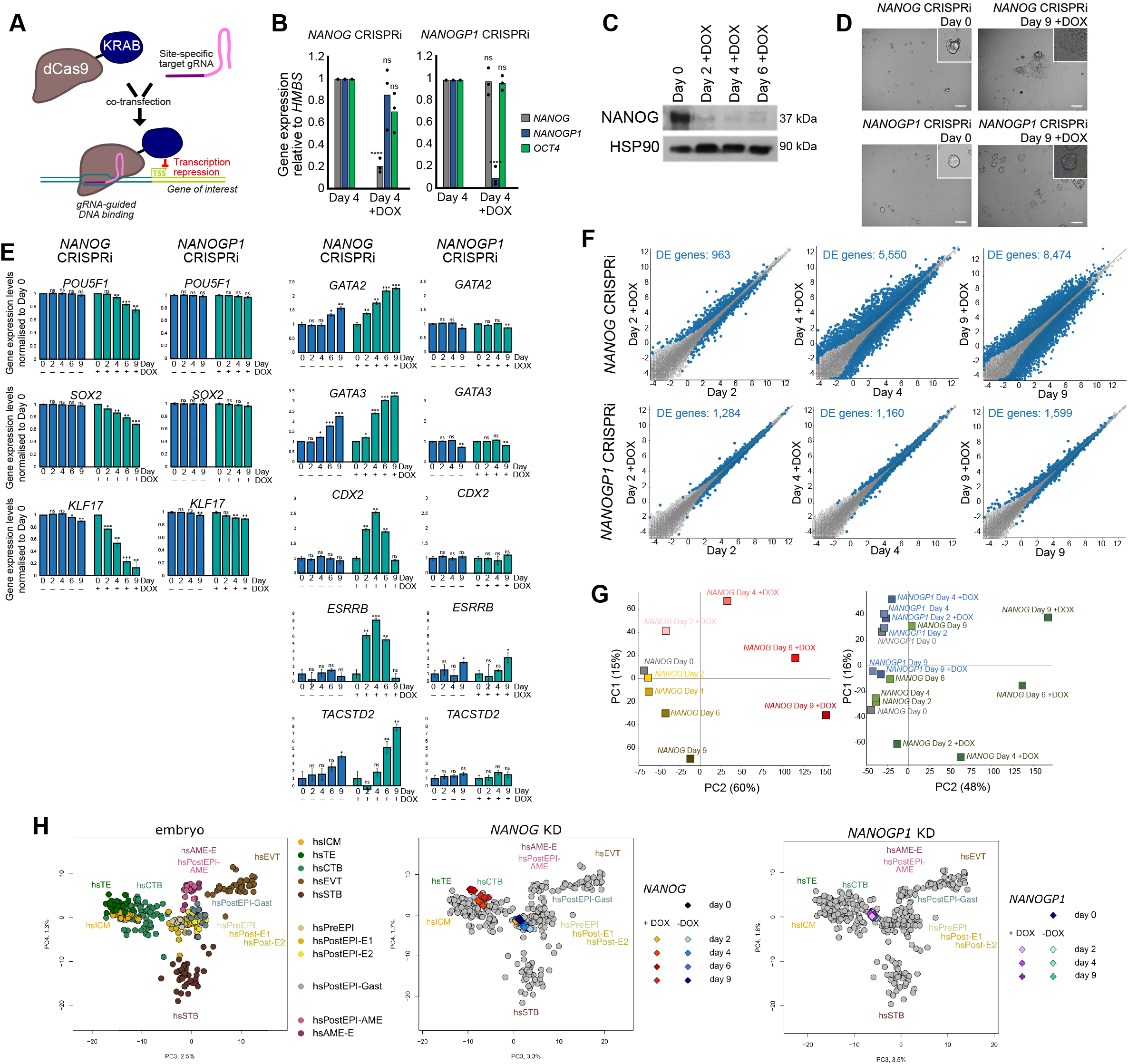
NANOG is required to maintain naïve pluripotency, but NANOGP1 is dispensable. **A)** DOX-inducible dCas9-KRAB CRISPRi to suppress *NANOG* and *NANOGP1* transcription in naïve hPSCs. **B)** CRISPRi knockdown of *NANOG* (left) and *NANOGP1* (right) in naïve hPSCs (t2iLGo medium). RT-qPCR values are relative to *HMBS* expression and normalised to the Day 4 samples. Mean and data points from three biologically independent samples. A t-test for each +/− DOX pair was performed (ns, not significant; p < 0.00005 (****)). **C)** Western blot shows reduced NANOG levels following DOX-induced *NANOG* CRISPRi in naïve hPSCs. **D)** Brightfield images of *NANOG* and *NANOGP1* CRISPRi naïve hPSCs on Day 0 and after 9 days of DOX treatment in t2iLGo medium. Inset images show representative colonies. Scale, 100 μm. **E)** Expression of undifferentiated (left) and trophectoderm markers (right) in *NANOG* and *NANOGP1* CRISPRi naïve hPSCs. Expression levels measured by RNA-seq are normalised to Day 0 samples. Data show mean from three biologically independent samples ± SD. A t-test with multiple testing correction was performed between each timepoint and the corresponding Day 0 sample (ns, not significant; p < 0.05 (*); p < 0.005 (**); p < 0.0005 (***)). **F)** Expression in *NANOG* (upper) and *NANOGP1* (lower) CRISPRi naïve hPSCs following DOX induction. Differentially expressed (DE) genes in blue (defined as p-adjusted < 0.05, Wald test). **G)** PCA plots show RNA-seq data of *NANOG* CRISPRi naïve hPSCs with and without DOX over a 9-day timecourse (left) and also with *NANOGP1* CRISPRi naïve hPSCs (right). Each data point is average of three independent samples. **H)** Left, PCA plot shows transcriptomes of annotated human embryo lineages (Xiang et al., 2020; Rostovskaya et al., 2022). On these maps, the transcriptomes of *NANOG* (centre) and *NANOGP1* (right) CRISPRi naïve hPSCs over a 9-day timecourse of DOX induction have been added. ICM, inner cell mass; TE, trophectoderm; CTB, cytotrophoblast; EVT, extravillous trophoblast; STB, syncytiotrophoblast; PreEPI, preimplantation epiblast; PostEPI, post-implantation epiblast; PostEPI-Gast, gastrulating stage; PostEPI-AME, post-implantation amniotic sac; AME, amniotic sac.

CRISPRi-mediated *NANOG* downregulation caused the naïve cells to lose their characteristic domed morphology and to visibly differentiate (Fig. 7D). Consistent with this, RNA-seq profiling over a 9-day time course revealed a strong transcriptional downregulation of naïve and core pluripotency factors (Fig. 7E). Transcriptionally upregulated genes were associated with the trophectoderm lineage, including *GATA2, GATA3, CDX2, ESRRB* and *TACSTD2*, and their induction was detected on day 2 and continued to increase in their expression up to day 9 (Fig. 7E).

In contrast, the downregulation of *NANOGP1* did not cause naïve hPSCs to induce the expression of trophectoderm marker genes or to change their morphology (Fig. 7D,E). Expression of pluripotent genes were unaltered (Fig. 7E) and, overall, far fewer differentially expressed genes were detected following *NANOGP1* downregulation compared to *NANOG* (Fig. 7F).

The transcriptional responses following the knockdown of *NANOG* or *NANOGP1* were distinct and well separated over the time course (Fig. 7G). Furthermore, by comparing the gene expression profiles to human embryo transcriptional data (Xiang et al., 2020), we further characterised the cell differentiation phenotype, and this also emphasised the differences following target gene depletion. *NANOG* knockdown naïve cells, starting from 4 days after doxycycline treatment, clustered with trophectoderm and cytotrophoblast cells of the embryo, whereas the earlier time-points (day 0 and day 2), non-induced cells, and all of the *NANOGP1* samples instead clustered closer to pre- and early post-implantation epiblast (Fig. 7H). These data confirm that *NANOG* is required to maintain naïve pluripotency, and establish that *NANOG*-depleted naïve hPSCs have similar transcriptional profiles to trophectoderm and cytotrophoblast lineages. In contrast to *NANOG*, the loss of *NANOGP1* expression does not disrupt the transcriptome of naïve pluripotent cells or cause trophectoderm differentiation. Additionally, *NANOGP1* did not provide functional redundancy for NANOG, as its expression was not sufficient to maintain naïve hPSCs in the absence of *NANOG*. In summary, these results demonstrated that downregulating the expression of *NANOG* in naïve hPSCs caused the loss of pluripotency, and that this function is not conserved for *NANOGP1*.

## DISCUSSION

To better understand the role of pseudogenes in human development and pluripotency, we characterised and studied the function of *NANOGP1*, a tandem duplicate of the transcription factor *NANOG*. We found that *NANOGP1* has overlapping but distinct expression patterns with *NANOG* in stem cell states and human embryo development. The restricted expression profile in epiblast, germ cells and hPSCs prompted us to investigate whether *NANOGP1* could have conserved functional activities in naïve pluripotency. First, we found that *NANOGP1* has the capacity for gene autorepression, as elevated expression of *NANOGP1* suppressed the expression of *NANOG* and *NANOGP1*. These findings additionally demonstrated that *NANOG* also has this function in human cells, which fulfils a prediction based on work in mouse pluripotent stem cells (Navarro et al., 2012). Second, *NANOGP1* was a strong inducer of naïve pluripotency when overexpressed in minimal reprogramming conditions, and was able to generate naïve hPSCs with comparable efficiencies to *NANOG*. These results are consistent with the ability of *NANOG* orthologues, and moreover the NANOG homeodomain by itself, to establish naive pluripotency in mouse (Theunissen et al., 2011). The intact homeodomain of *NANOGP1*, and the presence of NANOGP1 protein in human naive pluripotent cells, therefore provide elevated levels of an active form of the key pluripotency factor NANOG. Notably, we found that the homeodomain sequence of *NANOGP1* has been disabled in other primate species, including by a point mutation in *Rhesus macaque*, further supporting the likelihood that this domain has been conserved in human and other great apes. Lastly, because NANOG has dose-sensitive functions that are potentially mediated by concentration-dependent phase transitions (Choi et al., 2022), it is possible that NANOGP1 might contribute to these effects by lowering the critical concentration that is required for NANOG to form condensates.

Despite these functional capabilities, we also found that *NANOGP1* is not required to maintain naïve pluripotency *in vitro*. By engineering cells that expressed gene-specific CRISPR-interference to transcriptionally repress *NANOGP1*, we found that naïve hPSCs were unaffected by the robust knockdown of *NANOGP1*. Interestingly, the capacity of *NANOGP1* to induce naive pluripotency but is not required for its maintenance parallels another naive pluripotency factor – *KLF17* (Lea et al., 2021) In contrast, the knockdown of *NANOG* caused naive hPSCs to exit the naïve state and differentiate towards the trophoblast lineage, activating transcriptional programmes that matched trophoblast cells from human embryos. This finding demonstrates that, unlike mouse naïve pluripotent stem cells (Chambers et al., 2007; Novo et al., 2016), human naive cells require *NANOG*. It will be important to determine if this requirement is related to the specific capacity of human naïve cells to differentiate into trophoblast (Castel et al., 2020; Cinkornpumin et al., 2020; Dong et al., 2020; Guo et al., 2021; Io et al., 2021), which could underpin the different sensitivities to the loss of *NANOG*.

It is likely that the downregulation of *NANOGP1* has little effect in naive hPSCs because *NANOG* remains robustly expressed. However, we cannot rule out subtle effects including deficiencies following loss of *NANOGP1* that we have not yet identified. One interesting future direction would be to investigate whether the differences in predicted protein structures between NANOGP1 and NANOG create functional or regulatory differences. A prominent difference between the predicted NANOGP1 and NANOG proteins is a 39 amino acid deletion of the NANOGP1 N-terminus. The NANOG N-terminus has a role in transcriptional interference by attracting co-repressors of cell differentiation, thereby opposing the transactivation role that is mediated by the C-terminus (Chang et al., 2009). A key question, therefore, is whether NANOGP1 might lack this co-repression activity. The NANOG N-terminus is also a target for post-translational protein modifications, such as phosphorylation and ubiquitination, and the control of protein turnover (Oh et al., 2005). Future studies could therefore be aimed at determining whether there are differences in protein stability and perdurance between NANOG and NANOGP1, and, by implication, whether NANOGP1 might operate outside of the processes that act to control and limit NANOG activity.

Previous predictions based on mutation analysis proposed that *NANOGP1* is ~22 million years old (Booth and Holland, 2004). Our comparative phylogenetic analysis of primate genome assemblies suggests an older duplication date, of either approximately 40 Mya, between the divergence of apes and Old World monkeys (25-35 Mya) and the earlier divergence of New World monkeys (40-50 Mya), or still earlier before the divergence of New World monkeys from other primates. The availability and in some cases the quality of current primate genome assemblies is insufficient to distinguish between the two scenarios and this is a limitation of our study. More New World monkey and other primate genome assemblies would be informative, and also it was not possible in most cases to search for the informative ‘scars’ that might remain following *NANOGP1* duplication and deletion. Therefore, it is only possible at present to conclude that the duplication event took place at least ~40 Mya.

Our findings raise the question of why *NANOGP1* is retained in great apes but decayed in the genomes of lesser apes, Old World and New World monkeys. If NANOGP1 provides epiblast cells with higher levels of NANOG-like activity, then perhaps this relates to, and is informative to understand, the different developmental strategies between species. It is possible that the distinct modes of implantation (interstitial in great apes; superficial in New World and Old World monkeys), together with differences in the timing of blastocyst expansion and emergence of cell lineages, could point to a need to fine-tune transcription factor activities (Carter and Pijnenborg, 2011; Carter et al., 2015; Enders and Schlafke, 1986; Nakamura et al., 2016). To compare the functional role of transcription factors in early embryo development between different species, one future possibility could be to use stem cell-derived embryo-like models (Kagawa et al., 2022; Liu et al., 2021; Sozen et al., 2021; Yanagida et al., 2021; Yu et al., 2021) from different species as a representative and genetically tractable system.

The majority of duplications in the human genome are segmental duplications, which, in particular, are thought to drive evolution of great apes and humans (Marques-Bonet et al., 2009a; Marques-Bonet et al., 2009b). *NANOGP1*, however, was formed by tandem duplication, an older evolutionarily mechanism. Strikingly, a tandem duplication of *NANOG* has occurred and was conserved at least twice: once, forming *NANOGP1;* and once, at a substantially earlier point, forming *NANOGNB*, which has diverged to such an extent that was only recently recognised as a duplicate of *NANOG* (Dunwell and Holland, 2017). Independent *NANOG* duplications have also been reported in birds (Cañón et al., 2006), guinea pigs and some fish species (Scerbo et al., 2014). In all of these examples, the *NANOG* duplicates retain high similarity to their original ancestral sequences. These observations raise the possibility that the *NANOG*-containing region is somehow predisposed to duplication and retention of the duplication. In human, the chromosome region where *NANOG* is located also contains *DPPA3, OCT4P3* and another pluripotency factor *GDF3*, and collectively is called a ‘hotspot for teratocarcinoma’ due to the high rate of chromosomal abnormalities (Clark et al., 2004; Jong et al., 1990; Murty et al., 1990; Pain et al., 2005). Moreover, this region is also one of the most common amplification hotspots in hPSCs, which can accumulate large genomic duplications during hPSC culture (Adewumi et al., 2011). There may be relevant parallels between the seemingly beneficial amplification of the *NANOG*-containing region throughout evolution and the aberrant amplification of the region associated with cell adaptation. A study in yeast showed that genes that are highly expressed prior to duplication have a higher chance to be retained for a longer evolutionary period and in a wider phylogenetic range (Mattenberger et al., 2017). If highly transcribed genes are more likely to be duplicated and retained, this raises specific and important implications for the genetic control of early epiblast development, particularly as chromosome changes in these cells would be heritable.

Pseudogenes are defined as disabled or defective versions of protein-coding genes and have long been considered as non-functional elements. The majority of pseudogenes in the human genome are processed. However, there are over 2,000 unprocessed pseudogenes formed by duplication, many of which will have also copied their regulatory sequences. Careful annotation of pseudogenes, ideally supported by functional data, are important because they inform the reference list of genes and this impacts on whether sequence reads for the genes are mapped by default in genome assemblies or are included in genetic screens and other related methods. Here, CRISPR-based approaches to epitope tag an endogenous pseudogene, and to recruit transcriptional repressive machinery to the endogenous promoter, enabled us to selectively explore pseudogene function. By doing this, we established that *NANOGP1* is protein-coding and is expressed in pluripotent cells with functional activity. These results argue for the reclassification of *NANOGP1* to a protein-coding gene and that we should consider this factor as a gene, rather than a pseudogene. In addition to *NANOGP1*, we found other highly expressed pseudogenes of prominent pluripotency factors, such as *POU5F1* and *DPPA3*, and it is therefore important to investigate whether they too are protein-coding with functional properties. Defining pseudogene functionality and evolutionary conservation would help to uncover their involvement in species-specific developmental programmes and strategies.

## MATERIALS AND METHODS

### Human pluripotent stem cell lines

The use of human embryonic stem cells was carried out in accordance with approvals from the UK Stem Cell Bank Steering Committee. All cell lines used in this study were confirmed to be mycoplasma-negative. WA09/H9 primed hPSCs were obtained from WiCell (Thomson et al., 1998). WA09/H9 NK2 (Takashima et al., 2014) and chemically-reset WA09/H9 (Guo et al., 2017) naive hPSCs were kindly provided by Austin Smith (University of Exeter). The CRISPRi Gen1B primed hPSCs (Mandegar et al., 2016) were kindly provided by Bruce Conklin and Li Gan (Gladstone Institutes).

### Human pluripotent stem cell culture

All hPSC lines were maintained at 5% O_2_, 5% CO_2_ at 37°C in a humidified incubator. Naïve hPSCs were cultured in N2B27 media composed of 1:1 DMEM/F12 and Neurobasal, 0.5x B-27 supplement, 0.5x N-2 supplement, 2 mM L-Glutamine, 50 U/ml and 50 μg/ml penicillin-streptomycin and 0.1 mM β-mercaptoethanol (all ThermoFisher Scientific), supplemented either with 2 μM Gö6983 (Tocris), 1 μM PD0325901, 1 μM CHIR99021, and 20 ng/ml human LIF (all Wellcome-MRC Cambridge Stem Cell Institute) for t2iLGö medium (Takashima et al., 2014) or 1 μM PD0325901, 2 μM Gö6983, 20 ng/ml human LIF and 2 μM XAV939 (Cell Guidance Systems) for PXGL medium (Bredenkamp et al., 2019b; Rostovskaya, 2022; Rostovskaya et al., 2019). Naive hPSCs were grown either on irradiated MF1 mouse embryonic fibroblasts (MEFs) (Wellcome-MRC Cambridge Stem Cell Institute) on plates pre-coated with 0.1 %Gelatin (Sigma-Aldrich), or in feeder-free conditions using Geltrex Matrix (ThermoFisher Scientific) added to medium at a 1:300 dilution. Naïve hPSCs were passaged by 5 min incubation at 37 °C with Accutase (BioLegend). Primed hPSCs were cultured on plates pre-treated with 5 μg/ml Vitronectin (ThermoFisher Scientific) in mTeSR Plus medium (STEMCELL Technologies) and passaged by 5 min incubation at room temperature with 0.5 mM EDTA in PBS.

### *NANOGP1* epitope-tagging

CRISPR/Cas12a-mediated gene editing, described in (Zetsche et al., 2015), was adapted to epitope tag *NANOGP1*. Cas12a crRNA (IDT) targeting a region 10 bp upstream of the *NANOGP1* ATG site (5’-TGGGCCTGAAGAAAACCATCC-3’), and a repair template containing an epitope tag (V5 or 3xFLAG; Table S1), were designed using CRISPOR (http://crispor.tefor.net/). For cell nucleofection, 5.6 μg Alt-R A.s. Cas12a crRNA and 40 μg Alt-R A.s. Cas12a Ultra protein were pre-assembled for 15 min at room temperature, combined with 2 μl 200 pmol/ul repair template (all reagents produced by IDT) and transfected into cR-H9 naïve hPSCs using a Neon Transfection System (ThermoFisher Scientific). Each transfection reaction was performed using 1 million cells per 100 μl Neon Transfection tip and with 1300 V, 30 ms, 1 pulse settings. After transfection, the cells were transferred to PXGL naïve hPSC media supplemented with 10 μM Y-27632 (Cell Guidance Systems). To improve the rate of homology-directed repair, the cells were incubated in cold shock conditions (32°C) for 24 hr (Guo et al., 2018; Skarnes et al., 2019) at 5% O2, 5% CO2 in a humidified incubator. Additionally, 2 μM M3814 (DNA-dependent protein kinase inhibitor) (Sigma-Aldrich) was added to the cell media for 72 hr to repress non-homologous end joining DNA repair (Riesenberg et al., 2019). To improve survival, 10 μM Y-27632 was added to the cells for 2 h before cell transfection and was kept in the media for 72 h after the transfection. The resultant cR-H9 NANOGP1-tag cell lines were expanded in PXGL media.

### Inducible gene overexpression

To generate doxycycline-inducible gene overexpression vectors, gene cDNA was synthesised as a gBlocks Gene Fragment (IDT), cloned into a pCAG-IRES-Puro backbone vector (Niwa et al., 1991) and amplified with primers containing an *attB* sequence at their 5’ ends (Table S2). The amplification product (attB-gene cDNA-attB) was cloned into a TetON-GFP/RFP plasmid kindly provided by Andras Nagy (Woltjen et al., 2009) using a Gateway strategy (Hartley, 2003; Hartley et al., 2000) and was validated by Sanger sequencing (Genewiz). TetON plasmids, as well as plasmids encoding constitutively-expressed reverse tetracycline-regulated transactivator gene (pCAG-rtTa-Puro) and a piggyBac transposase (pCyL43) (Wang et al., 2008) were transfected into primed H9 hPSCs using an Amaxa 4D nucleofector (Lonza) with the setting CB-150. Stable cell lines were generated by 1 μg/ml puromycin selection for 48 hr, followed by transient gene induction by adding 1 μM doxycycline for 48 h and flow sorting for fluorescent reporter expression. For all assays that included more than one cell line, the same sorting gate was used to sort reporter-positive cells in order to establish lines with similar gene expression level.

### Primed to naïve hPSC chemical reprogramming

Primed TetON-NANOGP1-GFP H9 hPSCs were reprogrammed into the naïve state using a chemical reprogramming method (Guo et al., 2017; Rugg-Gunn, 2022). Feeder-free cultures of primed hPSCs were passaged onto feeders in mTeSR Plus medium supplemented with 10 μM Y-27632 at a density of 1×10^4^ per cm^2^ (Day 0) and provided with mTeSR Plus medium without Y-27632 on the following day. On Day 2, the medium was changed to chemical reprogramming medium 1 (cRM-1), composed of N2B27 medium supplemented with 1 μM PD0325901, 10 ng/ml human LIF and 1 mM valproic acid sodium salt (Sigma-Aldrich). Starting from Day 4, the medium was changed daily. On Day 5, cRM-1 medium was replaced with chemical reprogramming medium 2 (cRM-2), composed of N2B27 medium supplemented with 1 μM PD0325901, 10 ng/ml human LIF, 2 μM Gö6983 and 2 μM XAV939. After several passages, the culture became homogeneous and was transferred to t2iLGö medium.

### *NANOGP1-mediated* reprogramming

Primed H9 hPSC lines transfected with either *TetON-NANOGP1-GFP* (all three *NANOGP1* isoforms separately) plus *TetON-KLF2-RFP*, or with *TetON-NANOG-GFP* plus *TetON-KLF2-RFP*, were reprogrammed as described in (Takashima et al., 2014). Prior to reprogramming, primed hPSCs were treated with 1 μM doxycycline for 48 h and flow-sorted for GFP+ signal or GFP+/RFP+ double-positive signal to establish transgenic lines with the equivalent level of reporter expression. Transgenic lines were then plated on feeders in KSR/FGF2 medium comprising of 80 % Advanced DMEM, 20 % Knockout Serum Replacement (KSR), 2 mM L-Glutamine, 50 U/ml and 50 μg/ml Penicillin Streptomycin, 0.1 mM b-mercaptoethanol (all ThermoFisher Scientific), 4 ng/ml basic Fibroblast Growth Factor (Wellcome–MRC Cambridge Stem Cell Institute) supplemented with 10 μM Y-27632 (Day 0) and, on the following day, the medium was changed to KSR/FGF2 supplemented with 1 μM doxycycline. On Day 2, medium was changed to t2iL medium, composed of N2B27 medium with 1 μM PD0325901, 1 μM CHIR99021, 10 ng/ml human LIF, and supplemented with 1 μM doxycycline. t2iL medium was changed daily and cells were passaged every 5 days. On Day 12, doxycycline was withdrawn and 5 μM Gö6983 was added. Reprogrammed cells were propagated in t2iLGö medium on feeders.

### Inducible gene expression knockdown

dCas9-iKRAB Gen1B CRISPRi *NANOGP1* and CRISPRi *NANOG* hPSC lines were generated as follows. Gene-specific gRNA oligonucleotides were phospho-annealed and cloned into pgRNA-CKB (pCAG-mKate2-T2A-bsd) vector (Mandegar et al., 2016), pre-digested with BsmBI (NEB) and pre-treated with FastAP (ThermoFisher Scientific). The *NANOGP1* gRNA sequence was designed and validated in this study, and the *NANOG* gRNA sequence was from (Mandegar et al., 2016). Sequences are in Table S3. Linearised vector and phospho-annealed gRNA oligonucleotides were ligated at room temperature overnight with T4 DNA Ligase (ThermoFisher Scientific). Ligated products were validated by Sanger sequencing (Genewiz). Sequencing primers used were 5’-GAGATCCAGTTTGGTTAGTACCGGG-3’ and 5’-ATGCATGGCGGTAATACGGTTAT-3’.

CRISPRi Gen1B primed hPSCs (Mandegar et al., 2016) were nucleofected with the *NANOGP1* and *NANOG* gRNA plasmids using Amaxa 4D Nucleofector (setting CB-150), selected by blasticidin treatment (8 μg/ml for 5 days) and flow-sorted for mKate2 expression. Primed CRISPRi Gen1B *NANOGP1* and *NANOG* lines were reprogrammed into the naïve state using 5i/L/A-mediated resetting (Fischer et al., 2022; Theunissen et al., 2014). To do this, primed feeder-free cultures were passaged onto feeders in mTeSR Plus medium supplemented with 10 μM Y-27632 at a density of 2×10^4^ per cm^2^ (Day 0). On Day 1, mTeSR Plus was replaced with 5i/L/A medium composed of N2B27 medium supplemented with 1 μM PD0325901, 20 ng/ml human LIF and 20 ng/ml Activin A (Wellcome–MRC Cambridge Stem Cell Institute), 1 μM IM12, 0.5 μM SB590885, 10 μM Y-27632 and 1 μM WH-4-023 (all from Cell Guidance Systems). Cultures were passaged every 5 days and transferred to t2iLGö medium on Day 18. CRISPRi was induced with 1 μM doxycycline.

### Alkaline phosphatase activity

Colony formation assay was performed in combination with alkaline phosphatase (AP) staining (Štefková et al., 2015). Human PSCs were dissociated into single cells and plated into the experiment-specific medium onto feeders in 6-well plates. On Day 12, the cells were assayed for AP activity and imaged using a Zeiss Axio Observer Z1 with a 10X objective lens and Zeiss AxioVision software. Cells were fixed with 4% paraformaldehyde (PFA; Agar Scientific) in PBS, incubated in Alkaline Phosphatase staining solution (Merck) for 15 min and washed with PBS twice. The number of AP positive colonies was counted.

### Protein immunoprecipitation

All buffers used in this protocol were made with distilled water, were pre-chilled to 4°C, and contained cOmplete EDTA-free protease inhibitor. All centrifugation steps were performed at 4°C. NANOGP1-V5 and NANOGP1-3xFLAG hPSCs were harvested and centrifuged for 5 min at 300 x g, with 5×10^6^ cells per immunoprecipitation sample. To fractionate nuclei, pellets were resuspended in ice cold Buffer A (10 mM HEPES, 1.5 mM MgCl2, 10 mM KCl, 0.5 mM DTT, 0.05% NP40 and 250 u/ml Benzonase Nuclease (Sigma-Aldrich), incubated for 10 min on ice and centrifuged for 10 minutes at 2,000 x g. Cell pellets were resuspended in 376 μl Buffer B (5 mM HEPES, 1.5 mM MgCl2, 0.2 mM EDTA, 0.5 mM DTT, 26% Glycerol, and 250 u/ml Benzonase Nuclease, followed by 24 μl of 5 M NaCl. The resulting mix was homogenised using a Dounce on ice. Cell suspensions were kept on ice for 30 min followed by centrifugation for 20 min at 17,000 x g. The supernatant was analysed by Bradford assay and stored on ice. Using a magnetic rack, Protein A and Protein G Dynabeads (Thermofisher Scientific) were washed twice with IP dilution buffer (150 mM Tris-HCl pH 7.5, 150 mM NaCl, 0.5 mM EDTA). Then, 5 μg of anti-V5 and anti-FLAG antibodies (Table S4) were added to the Protein G and Protein A magnetic beads, respectively, which were diluted in 500 μl IP dilution buffer. Tubes were kept on a rotating wheel at 4°C overnight. Next day, the beads were washed three times in the IP dilution buffer. Then, 475 μg (95%) of the nuclear protein obtained in the lysis step was added to the beads. 25 μg (5%) of each protein sample were set aside as input. Immunoprecipitation samples were rotated at 4°C overnight. Next day, beads were resuspended in the IP dilution buffer and washed for a total of three washes. To elute the immunoprecipitated complexes, beads were resuspended in 20 μl 5x protein loading dye and boiled at 75° for 10 min. The eluate was diluted at 1x concentration, stored at −80°C and used in Western blot assays.

### Western blotting

Protein samples were extracted from frozen cell pellets, resuspended in ice-cold RIPA buffer (25 mM Tris/HCl, 140 mM NaCl, 1% Triton X-100, 0.5% SDS, 1 mM EDTA, 1 mM PMSF, 1 mM Na_3_VO_4_, 1 mM NaF) supplemented with cOmplete protease inhibitor (Roche, 1836170). Cells were lysed by incubating on ice for 30 minutes. Lysates were centrifuged at 16,000 x g for 30 min at 4°C. Protein concentration in supernatants was quantified using the Bradford assay. An appropriate volume of each lysate (containing 20-50 μg of the protein) was mixed with a 5x protein loading dye (5%β-mercaptoethanol, 0.02% bromophenol blue, 30% glycerol, 10% SDS, 250 mM Tris-Cl, pH 6.8), and incubated at 90°C for 5 min. Samples were vortexed and placed on ice. Protein samples were run on a polyacrylamide vertical gel and transferred onto a polyvinylidene fluoride (PVDF) membrane using iBlot gel transfer system. The membrane was blocked with 5% milk (Sigma-Aldrich) in TBST (Tris-buffered saline + 1% Tween20 (Sigma-Aldrich) for 1 hr at room temperature Primary antibody was applied in TBST + 5% milk overnight at 4°C. Next day, the membrane was washed three times with TBST and (HRP) conjugated secondary antibody was applied for 1 hr at room temperature. The membrane was washed three times and visualised by ECL or IRDye conjugated secondary antibodies. Antibody details are provided in Table S4.

### Immunofluorescence microscopy

Human PSCs were fixed in 12-well cell culture plates for 15 min at 4°C in 4 % PFA in PBS, washed once with PBS and permeabilised with 0.4 % Triton X-100 (Sigma-Aldrich) in PBS for 10 min at room temperature. Non-specific antibody binding was minimised by incubating cells with 3 % BSA (Sigma-Aldrich) + 0.1 % Triton X-100/PBS for 1 h at room temperature. The cells were incubated with the appropriate primary antibody in 3 % BSA + 0.1 % Triton X-100/PBS overnight at 4°C, before being washed four times with 0.1 % Triton X-100/PBS and incubated with the appropriate secondary antibodies in 3 % BSA + 0.1 % Triton X-100/PBS for 1 h at room temperature in the dark. Finally, the cells were washed three times in 0.1 % Triton X-100/PBS (for nuclei staining 1 μg/mL DAPI (Tocris) was added to the first wash) and two times in PBS. Wells were then filled with PBS, plates were sealed and stored at 4°C. Antibody details are provided in Table S5. Imaging was performed at the Babraham Institute Imaging Facility using a Nikon Live Cell Imager with a 20X objective lens.

### Flow cytometry

Cells were dissociated with Accutase, washed with 2 % FBS in PBS (Wash Buffer) and filtered through 50 μm sterile strainers (Sysmex). Antibody labelling was performed by incubating cells in a Brilliant Stain Buffer (BD Biosciences) with antibodies for 30 min at 4°C in the dark. This was followed by a wash in Wash Buffer, cell pelleting at 300 x g for 3 min and re-suspending the cells in 300 μl of the Wash Buffer. To identify live and dead cells, 0.1 μg/mL DAPI (Tocris) or Fixable Viability Dye eFluor 780 (eBioscience) was used. Antibody details are listed in Table S6. Flow cytometry analysis was performed on BD LSR-Fortessa at the Babraham Institute Flow Core. Cell sorting experiments were performed on BD Influx or BD FACSAria Fusion. Data processing and downstream analysis were performed using FlowJo V10.1.

### RNA-sequencing

RNA was extracted using an RNeasy Mini Kit (Qiagen). Indexed libraries were made using 0.5 μg RNA per sample with NEBNext Ultra™ RNA Library Prep Kit for Illumina with the Poly(A) mRNA Magnetic Isolation Module (NEB) and NEBNext Multiplex Oligos for Illumina (NEB). Agilent Bioanalyzer 2100 and KAPA Library Quantification Kit (KAPA Biosystems, KK4824) were used to identify library fragment size and concentration. Samples were sequenced as 75 bp single-end libraries on Illumina NextSeq 500 at the Babraham Institute Sequencing Facility, which generated 14-35 million uniquely mapped reads per library.

Sequencing files were analysed by FastQC v0.11.9 (https://www.bioinformatics.babraham.ac.uk/projects/fastqc/). RNA-sequencing reads were trimmed using Trim Galore v0.4.2 software (https://github.com/FelixKrueger/TrimGalore) to remove the adaptor sequences. Then, using HISAT2 v2.0.5 (Kim et al., 2015) guided by the Ensemble v70 gene models, trimmed reads were mapped to the human GRCh38 genome (Aken et al., 2016). Sequencing data was imported using Seqmonk software (http://www.bioinformatics.babraham.ac.uk/projects/seqmonk/). DESeq2 was used to identify genes expressed differentially (cut-off of p < 0.05 without independent filtering and after testing correction). To correct for the library size and variance among counts, regularised log transformation was applied prior to data visualisation. Principle component analysis (PCA) was performed using the top thousand most variable genes across the experiment, and the 1^st^ and 2^nd^ PCs were plotted.

### Polymerase chain reaction and genotyping primers

Polymerase chain reaction (PCR) was used to amplify various genomic and plasmid DNA fragments. PCR reactions were run in a BioRad Thermal Cycler T100. Polymerases Q5 HiFi (NEB), LongAmp Taq (NEB) and HotStarTaq (Qiagen) were used according to the manufacturer’s instructions. Primer sequences used in PCR reactions, genotyping and DNA Sanger sequencing can be found in Table S7.

### RT-qPCR

RNA was extracted using RNeasy Mini Kit (Qiagen) and then converted to cDNA using QuantiTect Reverse Transcription Kit (Qiagen). cDNA was diluted to 60 ng/μl and used in RT-qPCR using SYBR Green Jump Start Taq (Sigma-Aldrich) with 200 nM Forward and Reverse primers (Sigma-Aldrich; designed using Primer3 software (Untergasser et al., 2012). Samples were run in technical triplicates on 96-well plates on Bio-Rad CFX96 or 384-well plates on Bio-Rad CFX384. The results were analysed using the delta-delta cycle threshold method (relative quantity = 2^-ΔΔCt^) for which technical triplicates were averaged and normalised to the expression of a housekeeping gene *HMBS*. Data values represent Mean ± Standard Deviation of three biological replicates, unless stated otherwise. Statistical analyses are described in the figure legends. *NANOG* and *NANOGP1* expression in hPSCs was quantified using RT-qPCR primers, designed and validated to distinguish between the two genes. These two primer pairs, as well as other gene-specific primer sequences can be found in Table S8.

### Bioinformatics

#### Identification of *NANOGP1* transcript variants

To identify putative *NANOGP1* transcripts, a combination of in-house generated datasets of naïve hPSCs as well as publicly available data from (Theunissen et al., 2016) (GEO accession GSE84382), (Pastor et al., 2016) (GEO accession GSE76970) and (Takashima et al., 2014) (ENA accession PRJEB7132) was used. All raw data was processed with Trim Galore (adapter and quality trimming, v0.6.5) and mapped to the human GRCh38 genome using HISAT2 (v2.1.0; options --dta --sp 1000,1000), guided by known splice sites from Ensembl release 94 (Homo_sapiens.GRCh38.94.gtf).

To find evidence for splicing, aligned reads were first imported into SeqMonk (v1.43.1) as introns rather than exons, which effectively uses the CIGAR operation ‘N’ as the start and end coordinates of putative introns. Multi-mapping reads were filtered out (MAPQ >= 20).

To identify likely exons, reads were then imported into SeqMonk as standard i.e., spliced, RNA-seq reads (MAPQ >=20). Using read counts of exonic reads and introns identified as described above, the data was inspected and manually curated further to identify potential *NANOGP1* transcript variants. Transcript candidates appearing well supported by both exonic and intronic reads were termed *NANOGP1* isoform 1-3 and taken forward for further analyses. GTF/GFF files were generated for *NANOGP1* isoforms 1-3 and included as additional annotations for both HISAT2 mapping and further analyses in SeqMonk.

To identify potential open reading frames of *NANOGP1* isoforms 1-3 their hypothetical cDNA sequences were then screened for open reading frames (ORF) using the NCBI Open Reading Frame Finder tool (https://www.ncbi.nlm.nih.gov/orffinder/). The longest ORFs, resulting in predicted proteins between 255 and 266 amino acids in length, were taken forward for multiple sequence alignments (ClustalW) and additional analyses.

#### Disambiguation of *NANOG* and *NANOGP1*

To investigate the cross-mapping of reads from the *NANOG* to the *NANOGP1* locus, and vice versa, cDNA sequences for *NANOG* (NANOG-201, Ensembl) and *NANOGP1* (isoform 1) were used and converted to simulated FastQ files (as 43bp (like in Petropoulos et al., 2016) or 100bp single-end reads, in steps of 1bp from start to end). These *NANOG* and *NANOGP1* FastQ files were then aligned to the human GRCh38 genome (using HISAT2, v2.1.0); the amount of cross-mapping was either negligible or non-existent for unfiltered or multi-mapping filtered (MAPQ >=20) reads, respectively.

#### Human embryo data processing

The RNA-seq data of 1481 human embryo single cells from Petropoulos et al., 2016 were downloaded (accession number ERP012552) and categorised into the following groups: 8c, MOR, eICM, eTE, EPI, TE, PE, eUndef, Inter. Cell annotations were taken from Stirparo et al. 2018. The data were mapped to the human GRCh38 genome using HISAT2 (v2.1.0; options --dta --sp 1000,1000), guided by known splice sites from Ensembl release 94 (Homo_sapiens.GRCh38.94.gtf) to which a custom *NANOGP1* mRNA annotation had been added manually. Reads were then filtered for unique alignments (MAPQ > 20), and log2 RPM counts for genes were calculated with SeqMonk (v1.43.1; assuming non-strand specific libraries and merging transcript isoforms). Beanplots of expression values for genes of interest were then calculated for different developmental stages using the beanplot library in R (in RStudio).

The RNA-seq data of 557 human embryo single cells from (Xiang et al., 2020) were downloaded (accession number GSE136447) and categorised into the following groups: ICM, EPI, PrE, TrB. The data were mapped to the human GRCh38 genome using HISAT2 (v2.1.0; options --dta --sp 1000,1000), guided by known splice sites from Ensembl release 94 (Homo_sapiens.GRCh38.94.gtf) to which a custom *NANOGP1* mRNA annotation had been added manually. Reads were then filtered for unique alignments (MAPQ > 20), and log2 RPM counts for genes were calculated with SeqMonk (v1.43.1; assuming non-strand specific libraries and merging transcript isoforms). Violin plots of expression values for genes of interest were then calculated for different epiblast developmental stages using the ggplot2 package in R (in RStudio).

### Evolutionary genetics

To investigate the genomic structure of the *NANOG/NANOGP1* locus throughout evolution, the most recent assemblies of nine primate species (Table S9) were analysed. Approximate genomic coordinates of *NANOG* and *NANOGP1* (if present) were identified using BLAST (Basic Local Alignment Search Tool (BLAST)) and Needle (Madeira et al., 2019) pairwise sequence alignment tools. Within each assembly, a ~250 kilobase genomic region including *NANOG, NANOGP1* and their surrounding genes was extracted. The *NANOGP1* open reading frame for each species was also extracted. DNA and its corresponding amino acid sequences of *NANOG* and *NANOGP1* were aligned using MEGA (Tamura et al., 2007) and ClustalW (CLUSTAL W (improving the sensitivity of progressive multiple sequence alignment through sequence weighting, position-specific gap penalties and weight matrix choice), 2008). Codeml and codonml PAML (v4.8a) programs were run for the phylogenetic analysis of amino acid sequences with maximum likelihood under M0, M1, M7 and M8 models (Yang and Nielsen, 2000). Dotter (Barson and Griffiths, 2016) and Miropeats (Parsons, 1995) were used for visualising the *NANOG/NANOGP1* duplication site, detecting boundaries of the duplicated region and measuring conservation/divergence between the duplicated sequences since the duplication event.

The Gibbon nomLeu3.0 assembly was found to be not suitable for investigating the NANOG region due to having large gaps in the relevant region. To resolve this, unpublished gibbon genome assembly data based on long-read sequencing, kindly provided by Evan Eichler (University of Washington), was analysed. To visualise the *NANOG*-containing locus, human *NANOG* and *NANOGP1* sequence was mapped to gibbon contigs using Minimap2 (Li, 2018; Parsons, 1995).

For GC content calculation, enhancer regions were first extracted from human genome assembly (GRCh38 build) as FASTA files based on previously provided genomic coordinates. We then calculated GC content by dividing the sum of G and C nucleotide counts (G+C) to the total nucleotide count (G+C+T+A) at a genomic region. We used a 30 base-pair sliding-window approach to calculate GC content along the enhancer regions, and plotted GC percentages against genomic coordinates.

## Acknowledgements

We thank Austin Smith, Bruce Conklin and Li Gan for providing cell lines, and Evan Eichler for providing access to unpublished gibbon genome data. We are grateful to Paula Kokko-Gonzales and Amelia Edwards at the Babraham Institute Next Generation Sequencing Facility; Rachael Walker, Rebecca Roberts and team at the Babraham Institute Flow Core; and the Wellcome – MRC Cambridge Stem Cell Institute Tissue Culture Facility for providing reagents.

## Author contributions

Conceptualisation: K.M., P.J.R.-G.; Data curation: F.K.; Formal analysis: K.M., G.A., F.K., J.W., M.R., C.K., P.J.R.-G.; Funding acquisition: A.S., P.J.R.-G.; Investigation: K.M., G.A., J.W., M.R., A.B., S.W., P.J.R.-G.; Methodology: A.N.; Project administration: A.S., P.J.R.-G.; Supervision: A.S., P.J.R.-G.; Visualisation: K.M., G.A., J.W., M.R., P.J.R.-G.; Writing – original draft: K.M., P.J.R.-G.; Writing – review & editing: all authors.

## Data availability

RNA sequencing datasets have been deposited in the Gene Expression Omnibus (GEO) under the accession code of GSE204934.

## Supplementary Tables

**Table S1.**
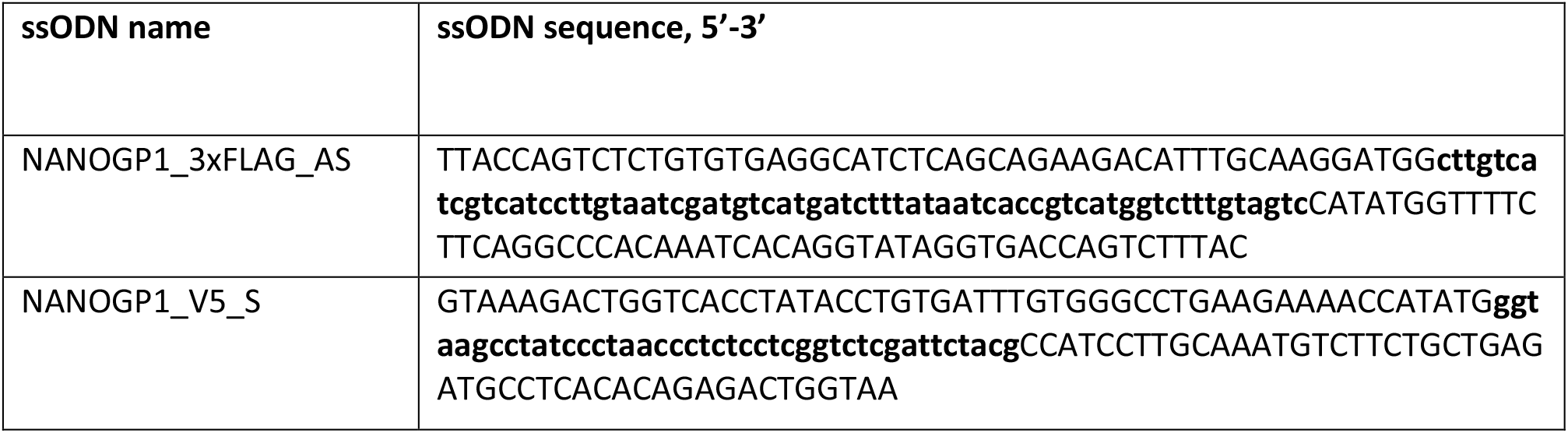
ssODN templates used in the *NANOGP1* epitope tagging experiment. AS – antisense strand. S – sense strand. Tag sequence is in bold. Homology arms are in capital letters.

**Table S2.**
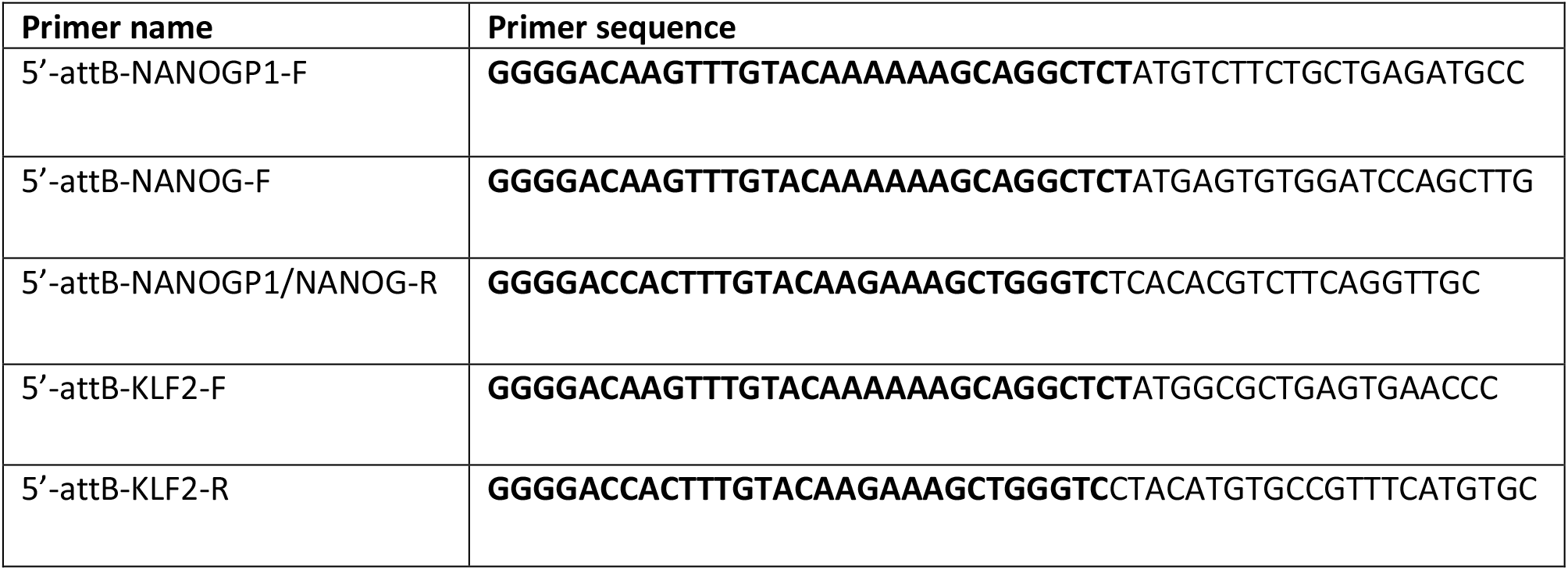
attB primer sequences used for generating TetON hPSC lines. attB sequences are in bold.

**Table S3.**
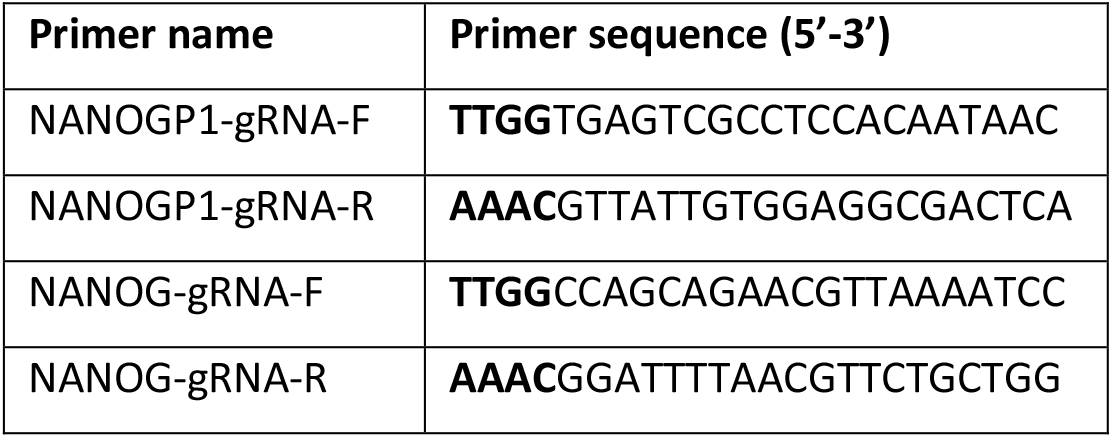
Primers designed for the pgRNA-CKB gRNA cloning. +/− values, distance from the gRNA PAM (Protospacer adjacent motif) site to the target gene transcription start site (TSS) in bp; ‘+’ indicates upstream location and ‘-’ indicates downstream location. ‘T’ and ‘NT’ indicate whether the gRNA targets the template or non-template strand, respectively. TTGG and AAAC in bold – overhangs added to clone phospho-annealed oligonucleotides to pgRNA-CKB using *BsmBI* restriction.

**Table S4.**
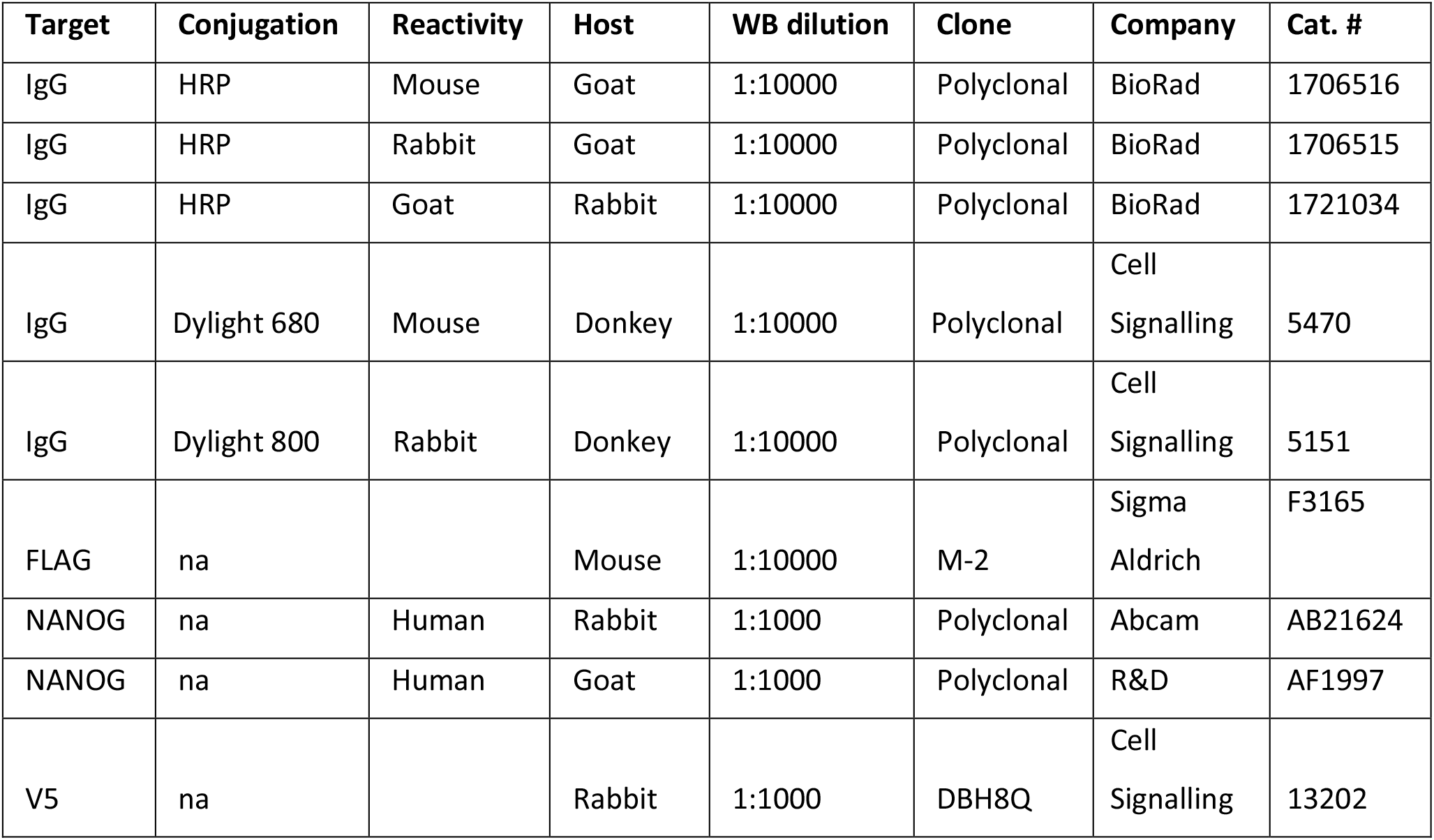
Western Blotting and protein immunoprecipitation antibodies. WB - Western Blotting. Na – not applicable.

**Table S5.** Immunofluorescent staining antibody details. CST - Cell Signalling Technology. SC – Santa Cruz. TFS - ThermoFisher Scientific. Na – not applicable.

| Target | Conjugate | Reactivity | Host | IF dilution | Clone | Company | Cat. # |
| --- | --- | --- | --- | --- | --- | --- | --- |
| IgG | AlexaFluor 555 | Goat | Donkey | 1:1000 | Polyclonal | TFS | A21432 |
| IgG | AlexaFluor 647 | Mouse | Donkey | 1:1000 | Polyclonal | TFS | A31571 |
| IgG | AlexaFluor 555 | Rabbit | Donkey | 1:1000 | Polyclonal | TFS | A31572 |
| NANOG | na | Human | Goat | 1:200 | Polyclonal | R&D | AF1997 |
| OCT4 | na | Human/mouse | Mouse | 1:300 | C-10 | SC | SC5279 |
| V5 | na | na | Rabbit | 1:150 | DBH8Q | CST | 13202 |

**Table S6.**
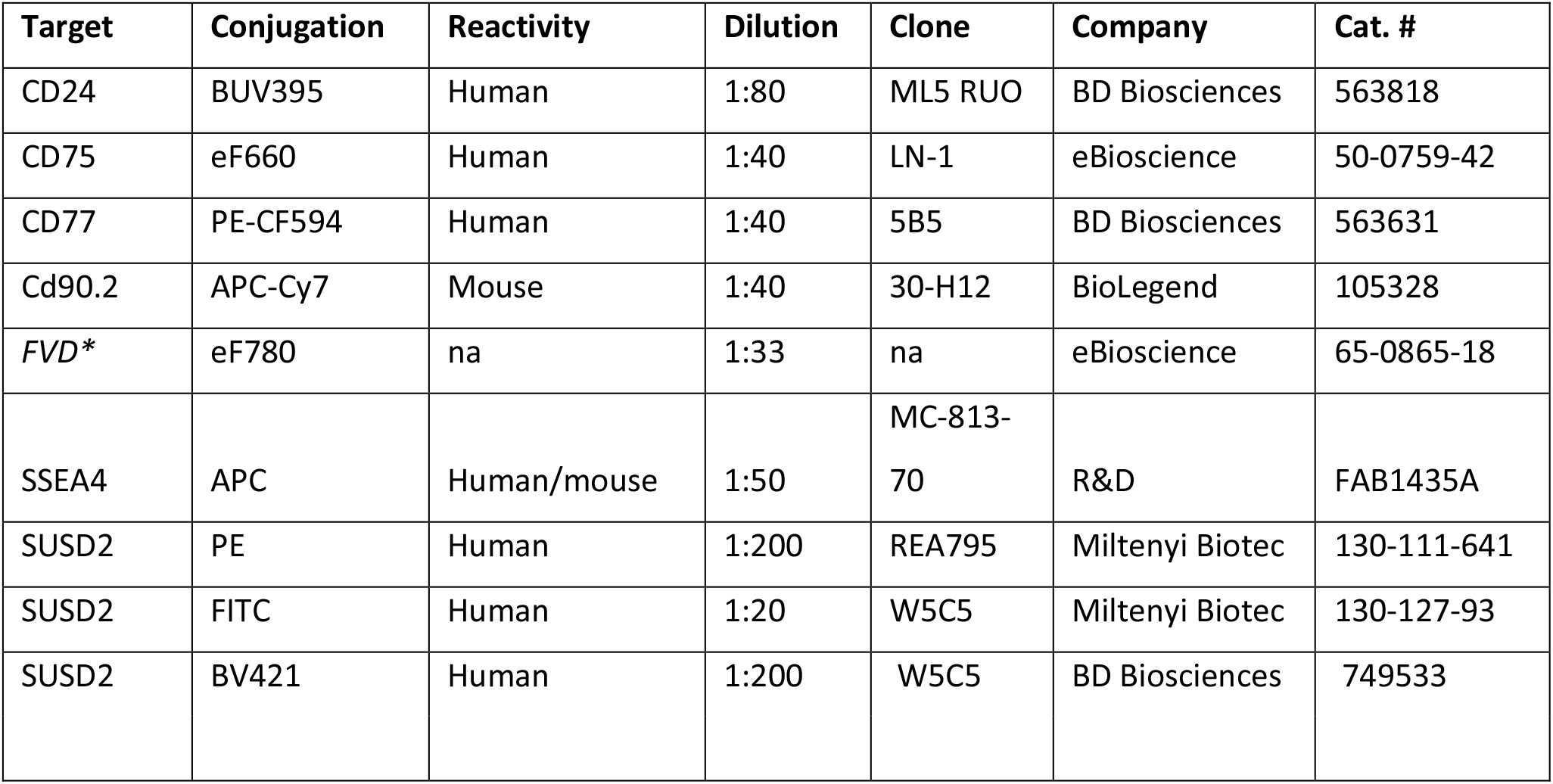
Flow cytometry antibodies. Dilution ratios per 100 μl buffer per 500,000 cells. FVD* - Fixable Viability Dye (not an antibody). Na – not applicable.

**Table S7.**
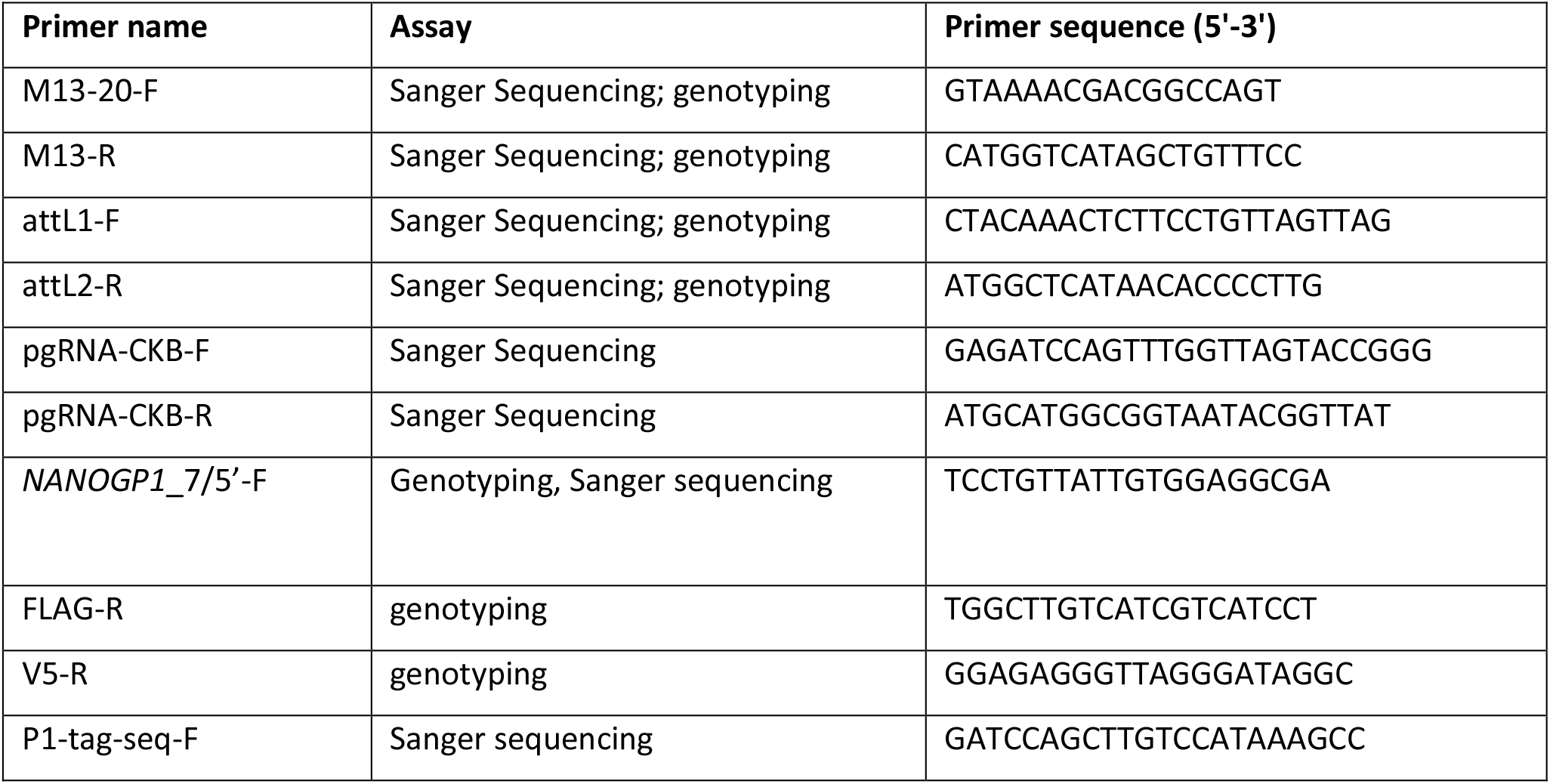
Primers used for genotyping, cloning validation and Sanger sequencing. F, R – forward and reverse primer orientation.

**Table S8.**
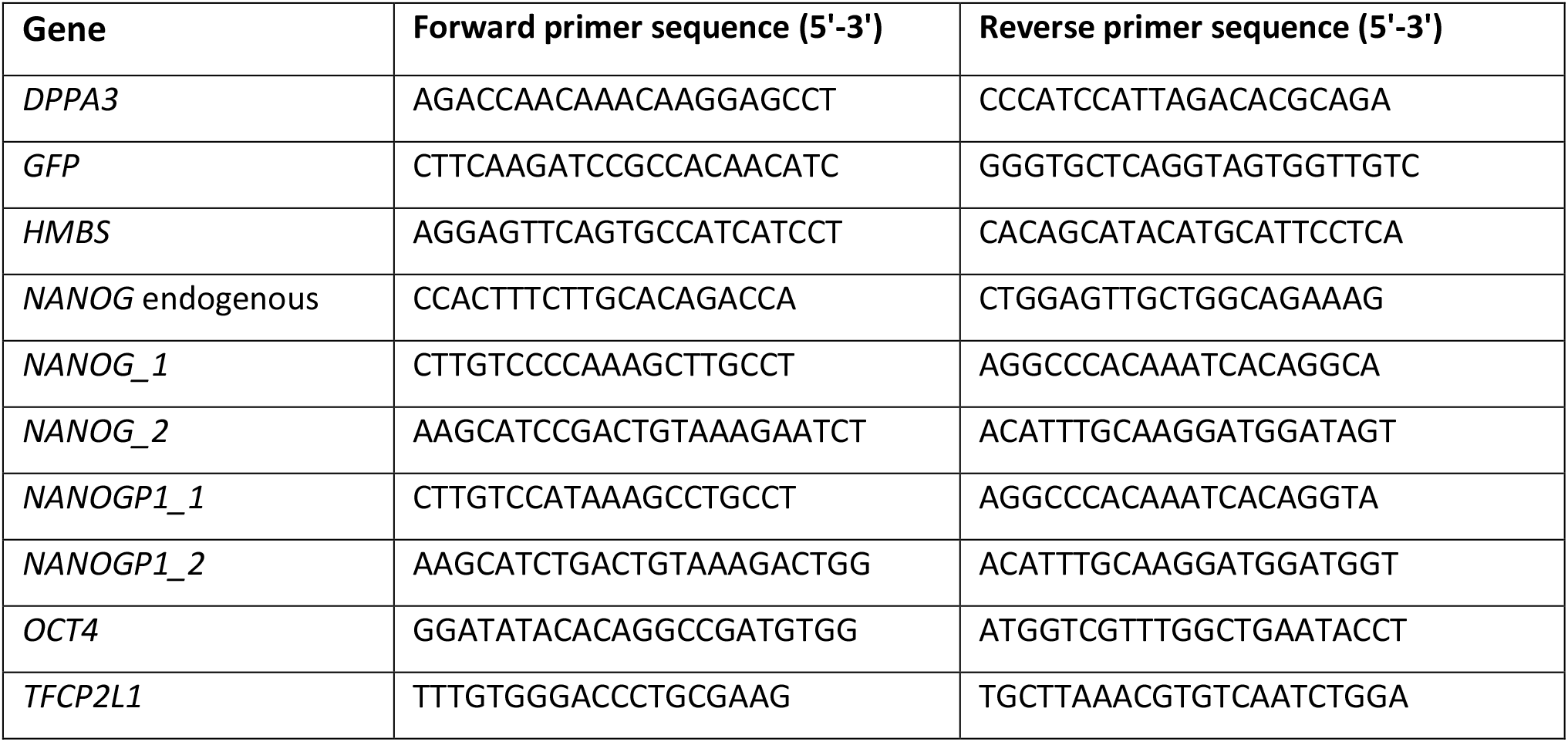
RT-qPCR primer sequences.

**Table S9.** Primate genome assemblies used in the evolutionary genetics assays.

| Species | Assembly | First release date |
| --- | --- | --- |
| Human | GRCh38 | 2013 |
| Chimpanzee | panTro6 | 2018 |
| Bonobo | panPan2 | 2015 |
| Gorilla | gorGor5 | 2016 |
| Orangutan | ponAbe3 | 2018 |
| *Gibbon* | *nomLeu3* | 2012 |
| Crab-eating macaque | macFas5 | 2013 |
| Rhesus macaque | rheMac8 | 2015 |
| Marmoset | calJac3 | 2009 |

